# Larval and adult traits coevolve in response to coastal oceanography to shape marine dispersal kernels

**DOI:** 10.1101/2022.12.19.521131

**Authors:** James H. Peniston, Scott C. Burgess

## Abstract

Dispersal emerges as an outcome of organismal traits and external forcings. However, it remains unclear how the emergent dispersal kernel evolves as a by-product of selection on the underlying traits. This question is particularly compelling in coastal marine systems where dispersal is tied to development and reproduction, and where directional currents bias larval dispersal downstream causing selection for retention. We modelled the dynamics of a metapopulation along a finite coastline using an integral projection model and adaptive dynamics to understand how asymmetric coastal currents influence the evolution of larval (pelagic larval duration) and adult (spawning frequency) life history traits, which indirectly shape the evolution of marine dispersal kernels. Selection induced by alongshore currents favors the release of larvae over multiple time periods, allowing long pelagic larval durations and long-distance dispersal to be maintained in marine life cycles in situations where they were previously predicted to be selected against. Two evolutionary stable strategies emerged: one with a long pelagic larval duration and many spawning events resulting in a dispersal kernel with a larger mean and variance, and another with a short pelagic larval duration and few spawning events resulting in a dispersal kernel with a smaller mean and variance. Our theory shows how coastal ocean flows are important agents of selection that can generate multiple, often co-occurring, evolutionary outcomes for marine life history traits that affect dispersal.

## Introduction

Dispersal has fundamentally important consequences for the demographic and genetic structure of populations, and how species respond and adapt to changing conditions (Clobert et al. 2012; Travis et al. 2013). As a result, it is important to understand how dispersal evolves. Decades of work by theoreticians has focused on dispersal propensity or dispersal distance as the evolving trait and shown how kin competition, inbreeding, and spatio-temporal variation can select for dispersal (Starrfelt and Kokko 2012). However, dispersal in nature arises as an outcome of the biological traits of organisms and external forcings (e.g., wind and water currents) that both affect movement, fitness, and the final distribution of dispersal distances (Burgess et al. 2016). Therefore, challenges remain in explaining how dispersal actually evolves in nature, rather than how it can evolve. That is, there is a need to identify which traits cause dispersal outcomes, and what factors cause selection on those traits to influence the pattern of dispersal that emerges and changes through evolutionary processes (Burgess et al. 2016). This challenge is particularly prevalent in marine systems, where dispersal is tied to early development in complex life cycles, and traits that influence dispersal outcomes are also traits that influence development and reproduction (e.g., egg size; Pringle et al. 2014).

In many marine invertebrates and fishes, adults are sessile or demersal, but their microscopic larval offspring are capable of dispersing great distances in ocean currents (kilometers to 100’s of kilometers in some species), mostly during obligate periods of development when larvae feed and are incapable of settling (Kinlan and Gaines 2003; Shanks 2009). However, the ease of larval dispersal in ocean currents creates problems for locating suitable settlement habitat after development. Along many coastlines, the average current is unidirectional over the timescales that dispersal occurs (Davis 1985). As a result, passive larvae drift downstream, which results in larvae being constantly washed away from settlement habitat (Gaylord and Gaines 2000; Largier 2003; Siegel et al. 2003). If there is not enough upstream retention, downstream dispersal ultimately leads to population extinction (Byers and Pringle 2006). This results in a ‘drift-paradox’, where adult populations persist despite the threat of a net downstream loss of larvae (Müller 1982; Speirs and Gurney 2001; Müller 1982; Speirs and Gurney 2001; Pachepsky et al. 2005; Shanks and Eckert 2005; Byers and Pringle 2006). Therefore, the ubiquity of alongshore currents in coastal habitats is expected to select for dispersal traits that increase upstream retention, but may also result in downstream dispersal as a consequence.

One trait that can influence dispersal and upstream retention is pelagic larval duration (Grantham et al. 2003; Shanks 2009; Treml et al. 2015; Cecino and Treml 2021). Shorter pelagic larval durations decrease the risk that passively dispersing larvae are transported and lost downstream on average (Siegel et al. 2003; Byers and Pringle 2006). Recent analyses considering the role of ubiquitous alongshore currents in coastal habitats have predicted that stronger currents should lead to the loss of pelagic larvae from marine life cycles all together (Pringle et al. 2014), suggesting that species with feeding larvae (planktotrophy) should only be found where currents are relatively weak. When mean currents are weak relative to the stochastic variation in current speed and direction, there are potential advantages to longer larval durations that relate to greater growth and survival in pelagic versus benthic habitats, but not necessarily for the dispersal they facilitate (Burgess et al. 2016; Meyer et al. 2021*a*; Iwasa et al. 2022). There is a large literature on the evolution of marine reproductive strategies based on egg size-number trade-offs where egg size affects larval development times depending on whether larvae feed or not (Vance 1973; Strathmann 1985; Emlet et al. 1987; Levitan 2000; Marshall and Keough 2007). This theory predicts that longer larval durations evolve when selection favors the production of many small offspring that feed for themselves away from adult habitat, but require longer to feed and develop independently to a size and stage required for settlement back into adult habitats (Strathmann 1974, 1990; Jackson and Strathmann 1981; Palmer and Strathmann 1981). Therefore, because egg size affects development time, which in turn affects the potential for upstream retention, ocean currents are expected to strongly modify how marine egg size-number trade-offs evolve (Reitzel et al. 2004; Shanks and Eckert 2005; Pringle et al. 2014; Álvarez-Noriega et al. 2020).

Despite most analyses on the causes of marine dispersal focusing on the traits of larvae, especially larval behaviors (Leis 2006; Morgan 2014), traits that affect dispersal and upstream retention may also include those of the less mobile adult stages. Parents not only control larval duration via the effects of egg size, but also the timing, frequency, and, in some cases, location in which offspring are released into coastal flow fields (Strathmann 1982; Morgan and Christy 1995; Reitzel et al. 2004). In particular, unidirectional alongshore currents often reverse direction on many coastlines due to wind or seasons. Releasing offspring on multiple occasions can increase retention by increasing the variability in advection that batches of larvae encounter among different releases. Accessing greater variability in currents over multiple releases increases the chance that enough of a parent’s lifetime reproductive output occasionally moves upstream against the average downstream flow compared to releasing only one batch of larvae (Byers and Pringle 2006). So while larval behaviors can also increase retention (Paris and Cowen 2004; Metaxas and Saunders 2009; Bottesch et al. 2016; Morgan et al. 2021; Burgess et al. 2022), adult traits also control dispersal by when and how often larvae are released into the current. Small dispersing larval stages and marine life histories are therefore not at the whim of strong physical forcing. Instead, the physical forcing itself causes selection on life history traits, and the pattern of dispersal that emerges can evolve (Burgess et al. 2016).

Our goal here was to develop theory that illustrates how coastal oceanographic processes affect the evolutionary outcome of traits that affect dispersal. Most previous theory has considered reproductive strategies in the absence of oceanography (Vance 1973; Levitan 2000), considered dispersal itself as the evolving trait rather than the underlying traits of the individuals that interact with currents to give rise to dispersal patterns (Shaw et al. 2019), or only focused on larval traits (Pringle et al. 2014). We use an adaptive dynamics framework to present new theory showing how asymmetric coastal currents influence the coevolution of pelagic larval duration and adult spawning frequency in coastal ecosystems. We consider lifetime dispersal kernels as the dispersal kernel of all larvae released over an individual’s lifespan. Our model shows how the evolutionarily stable combination of pelagic larval duration and spawning frequency changes with oceanographic conditions and indirectly affects the expectation for marine dispersal kernels. We show that for many realistic coastal oceanographic conditions, there are two evolutionarily stable life history strategies: one with a longer pelagic larval duration and higher spawning frequency and another with a shorter pelagic larval duration and lower spawning frequency, leading to different expected dispersal kernels under the same flow regime.

### Model Description

We model the dynamics of a metapopulation along a finite coastline using an integral projection model structured by space and age. The integral projection model framework is quite general and can be adapted for different assumptions by replacing any of the functions below with another suitable function.

We study the phenotypic evolution of pelagic larval duration (*T*_PLD_) and number of spawning events per individual parent (*N*_spawn_) by analyzing this model with an adaptive dynamics methodology, which allows for both frequency- and density-dependent dynamics (McGill and Brown 2007; Rees and Ellner 2016). Adaptive dynamics works by introducing a mutant with a small change in either *T*_PLD_ or *N*_spawn_ into a stabilized population of residents and calculating the invasion growth rate of its lineage. We determine the evolutionarily stable phenotypes by iterating this process and finding the phenotypes that cannot be invaded by mutants with small changes. Trait values that cannot be invaded are referred to as evolutionarily stable strategies (ESSs), and can be thought of as the endpoint, or outcome, of evolution. All evolutionarily stable strategies presented in our results were also convergence stable, and were thus what are referred to as continuously stable strategies (Brännström et al. 2013). In cases where there were two ESSs, we identified intermediate unstable equilibria by iteratively changing the initial trait values and then introducing mutants as described above.

In our results, we first discuss the evolution of *T*_PLD_ and *N*_spawn_ separately, that is, assuming one trait evolves while the other trait does not evolve. These analyses provide a useful understanding of how each trait independently affects each other’s evolution given the oceanographic currents. We then consider a model where both traits coevolve (*i.e*., affect each other simultaneously and reciprocally) and determine the evolutionarily stable combinations of trait values.

The ratio of mean alongshore flow (*U*) to short timescale fluctuations in currents (*σ*, that is, current fluctuations over the timescales captured by *U*) is a key descriptor of how oceanographic conditions affect dispersal evolution (Pringle et al. 2014). We will refer to this ratio as “scaled alongshore flow” and present results of life history evolution for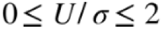, which captures a realistic range of oceanographic conditions (Robinson and Brink 2006).

#### Relationship between pelagic larval duration and fecundity

There is a trade-off between fecundity and egg size (*i.e.*, the more eggs an individual produces, the smaller each egg is). We model this size-number trade-off by defining the number of eggs released *f* as the total amount of material contributed to egg production *C* divided by the egg volume *s*_egg,_ that is 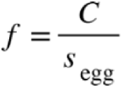.

There is empirical evidence that egg size affects pelagic larval duration depending on whether larvae feed or not. For feeding (planktotrophic) larvae, pelagic larval duration decreases with egg size, especially in echinoderms (Vance 1973; Emlet et al. 1987; Levitan 2000; Marshall and Keough 2007; Marshall et al. 2018). Here, we focus on feeding larvae because we are initially interested in explaining how long-distance dispersal is maintained in marine invertebrate and fish life histories, and to also compare our results to previous models of pelagic larval duration in coastal oceans (e.g., Pringle et al. 2014). However, future studies should consider non-feeding larvae, which generally have shorter pelagic larval durations, or other traits that affect larval duration.

We model the negative relationship between the pelagic larval duration and egg size following Pringle et al. (2014) such that

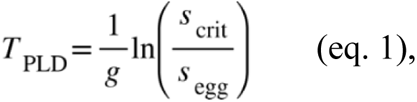

which can be rearranged as

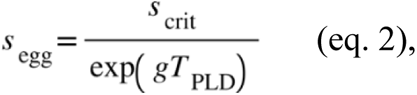

where *T_PLD_* is pelagic larval duration, *g* is the rate at which larvae gain mass during feeding, and *s*_crit_ is critical size that the egg must reach to settle. A given increase in egg size for smaller eggs reduces *T_PLD_* more than the same increase in egg size for larger eggs.

#### Larval mortality

We assume that the probability of larvae surviving through the pelagic stage decreases with pelagic larval duration and increases with egg size, such that it is given by

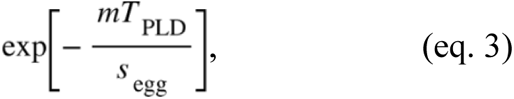

where *m* is the baseline daily larval mortality rate. The survival benefits of larger eggs can arise from larger larvae acquiring more energy, using proportionally less energy, or being less susceptible to predation (Marshall et al. 2018). More generally, equation (3) can be treated as a heuristic assumption that leads to a hump-shaped relationship between egg size and number of larvae that survive the pelagic phase. Such a relationship is an important concept in the marine life history literature because it allows for an optimal intermediate egg size, depending on the parameter values (Smith and Fretwell 1974; Levitan 2000).

Given equation 3 and the egg size-number tradeoff above, the number of larvae that survive the pelagic phase is

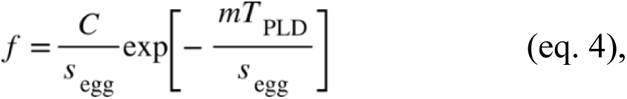

(Figure 1A shows eq. [4]). We only consider scenarios where larval growth rate is greater than the larval mortality rate (*g*-*m* > 0), because otherwise there is always selection for no pelagic larval stage and this has been explored previously (Pringle et al. 2014; Iwasa et al. 2022).

**Figure 1.**
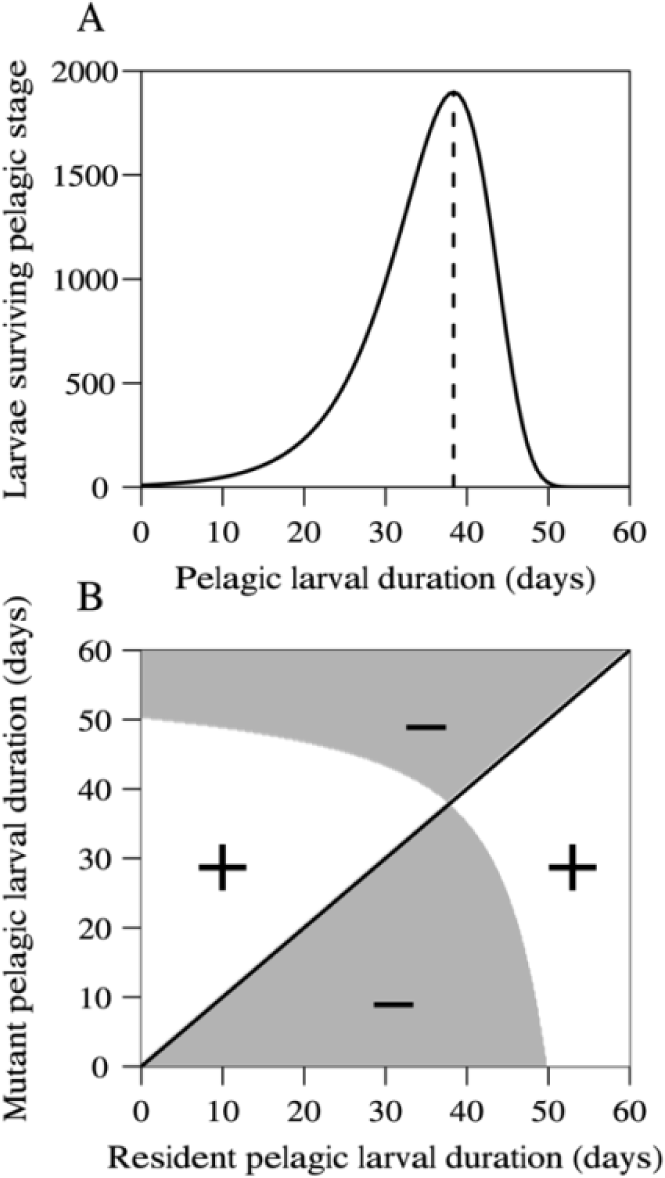
Evolution of pelagic larval duration (*T*_PLD_) without considering oceanography. Panel A: total number of larvae that survive the pelagic phase (*f*) as a function of pelagic larval duration given by equation 4. Dashed vertical line denotes the optimal pelagic larval duration that gives the maximum number of surviving larvae. Panel B: pairwise invasibility plot. Pairwise invasibility plots are a way of visualizing which mutant populations can invade which resident populations. Evolutionarily stable strategies occur at resident trait values that cannot be invaded by small mutations. In panel B, invasion fitness of the mutant is greater than 1 in the white areas and less than 1 in the grey areas. The black line denotes the 1:1 line where the invasion fitness of the mutant is 1. Note that there is only one evolutionary stable pelagic larval duration and it occurs at the value that maximizes the number of surviving larvae. Parameters: *τ*=4, C=0.1, *s*_crit_=0.0103, *g* = 0.16, *m* =5×10^-7^, *A_m_*=0.1.

#### Relationship between pelagic larval duration and the dispersal kernel

Dispersal is dependent on mean alongshore flow (*U*), the standard deviation in alongshore flow (*σ*), the Lagrangian decorrelation timescale (*τ*), and pelagic larval duration (*T*_PLD_). Siegel et al. (2003) show the mean dispersal distance (advection) is

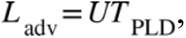

and the standard deviation in dispersal distance (diffusion) is

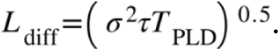

The term *σ*^2^*τ* is known as the eddy diffusion coefficient (Siegel et al. 2003). Based on basic oceanographic principles, a Gaussian dispersal kernel is used for passively dispersed larvae (Largier 2003). Thus, the probability density function describing the dispersal from location *x* to *y* (the dispersal kernel) is given by

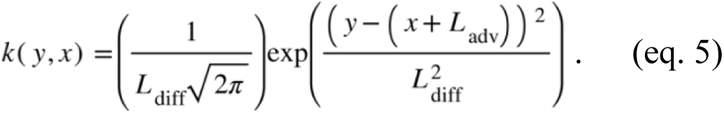

For a discussion of the sensitivity of upstream retention and selection on long pelagic larval durations to non-Gaussian dispersal kernels see (Pringle et al. 2009, 2014).

#### Recruitment competition

We assume that larvae can only settle at a site if there is an available microsite (e.g., space on a rock) and that there are only *K* microsites available at each site. Larvae have lottery competition for microsites (Chesson and Warner 1981; Warner and Chesson 1985). That is, the number of lineage *i* (mutant or resident) larvae that successfully settle is proportional to the relative frequency of that lineage among larvae arriving at the site. Therefore, the expected number of lineage *i* individuals that establish themselves at site *x* is given by

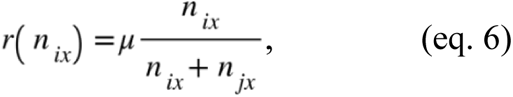

where *n_ix_* is number of lineage *i* larvae arriving at site *x* and *μ* is the number of unoccupied microsites, which is given by *K* minus the total number of adults that died the previous time step.

#### Post-settlement dynamics

After settlement, individuals live for up to *N*_spawn_ time steps and release larvae each time step. The probability of post-settlement individuals surviving to the next time step is given by the function *s*(*a*), where *a* is age. *s*(*a*) can be any age-dependent mortality function, but we will assume that all post-settlement individuals have the same mortality rate *A_m_* until reaching age *N*_spawn_, at which point they all die. For all presented results we divide the annual amount of material contributed to egg production *C* by *N*_spawn_ such that lifetime investment in egg production is fixed if there is no adult mortality (e.g., individuals could either produce 100 eggs ten separate times, produce 1000 eggs all during one spawning event, or anywhere along this range). We make this assumption because we are interested in the specific effects of spawning frequency *per se* and this assumption controls for the increased lifetime fecundity that might accrue with increased spawning frequency, or the amount of material contributed to egg production *C* that might increase as adults grow (Marshall et al. 2022). For comparison, we also ran analyses in which we allowed lifetime fecundity to increase with increased spawning frequency (*i.e.*, *C* was the same for all spawning events regardless of *N*_spawn_). In these analyses, *N*_spawn_ always evolved to the maximum possible value because there were only benefits to increasing *N*_spawn_. *T_PLD_* evolved as it would independently if *N*_spawn_ was fixed at that maximum value (Figure A1). Future studies could consider additional complexities that emerge from specific relationships between spawning frequency and fecundity.

#### Temporal fluctuations in alongshore flow

We investigate the effect of variation in alongshore flow rates that takes place on time scales equivalent to spawning frequency. For simplicity we will refer to these fluctuations as interannual variation in flow, which is accurate if time steps in our model are treated as years. However, the time steps can be treated as any unit of time longer than the Lagrangian decorrelation timescale (which we set to 4 days, Davis 1985), and “interannual” fluctuations can be interpreted as among time step fluctuations.

Interannual fluctuations in flow rates do not affect an individual’s lifetime dispersal kernel if they only spawn once, but when *N*_spawn_>1, we incorporate such interannual variation in flow rates into our model by replacing diffusion (*L*_diff_) with an estimate for the standard deviation of larval dispersal distance for all larvae released over the lifetime of an adult, *L*_diffeffect_, which was developed by Byers and Pringle (2006)

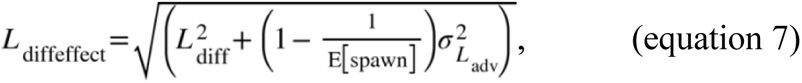

Where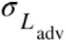 is the interannual standard deviation in *L*_adv_ and 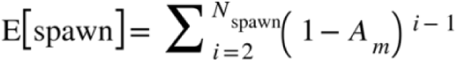 is an individual’s expected number of spawning events. Given our equation for *L*_adv_ above, 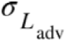 is *σ*_IA_*T*_PLD_, where 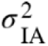 is the interannual variation in mean flow rates *U*. Note that, if *σ*_IA_ > 0 and *N*_spawn_ >1, then *L*_diffeffect_ > *L*_diff_. We checked the robustness of using this estimation of *L*_diffeffect_ using the direct stochastic simulation methods described in the Appendix.

#### Integral projection model

Metapopulation dynamics are given by the set of coupled equations

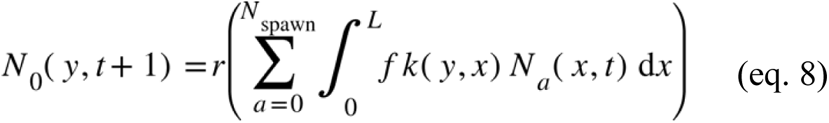

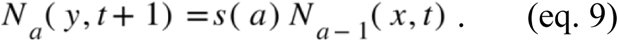

*N_0_*(*y*, *t*+1) is the number of larvae (age 0) individuals that settle at site *y* in timestep *t* + 1. *N_a_*(*y*, *t*) is the number of age *a* individuals at site *y* at timestep *t*. *f* is the number of larvae that survive the pelagic phase (eq [4]), *k(y,x)* is the dispersal kernel (eq [5]). *s*(*a*) gives the probability of age *a* individuals surviving to age *a*+1. *L* is the length of the coastline. If necessary for the study system, equations [8–9] can be adapted such that mortality rate, fecundity, and larvae size can vary with age.

#### Calculating invasion fitness

Invasion fitness *λ* is calculated as the initial growth rate of a mutant lineage introduced into, and competing with, a stabilized resident metapopulation. As long as the resident population’s local retention is positive, the resident population will fill all microsites along the coast. Therefore, the long-term stable resident distribution is *K* at all sites. The stable age distribution of residents is then given by (Dewi and Chesson 2003)

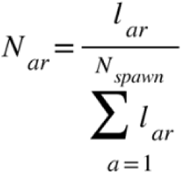

where *l_ar_* is the probability of surviving to age *a*. We can then rewrite the recruitment function (eq. 6) as

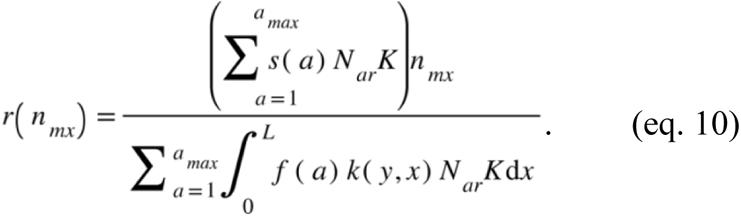

Note that the number of unoccupied sites *μ* is now given by the sum of age-specific adult mortality rates *s*(*a*) multiplied by the stabilized number of resident adults in that age class *N_ar_K*. Also, the denominator no longer includes mutants because invader density is by definition very low and can therefore be assumed to not affect density dependence. In essence, this is now a density-independent model where the resident metapopulation distribution is treated like an environmental factor affecting recruitment and growth of the mutant population.

Following Ellner and Rees (2006), we calculate invasion growth rates by numerically evaluating our model, with space discretized using the midpoint rule. Details of this protocol are included in the Appendix. We checked the robustness of our assumption that the resident population is stationary and our estimate of *L*_diffeffect_ by comparing our results to stochastic simulations which tracked the population dynamics of residents and stochastically varied flow rates. These simulations confirmed the key patterns of selection seen in our deterministic model (details in the Appendix). We focus our analysis on the deterministic model, however, because results of the stochastic model are dependent on specific assumptions about how stochasticity and density dependence are implemented, and while important processes could emerge from the stochasticity, the goal of this paper is to make general conclusions about the evolution of *T*_PLD_ and *N*_spawn_ in coastal systems.

#### Default parameter choices

We evaluated values of *U*, *σ*, 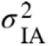, and *τ* that matched realistic ranges of coastal oceanographic conditions (Davis 1985; Lumpkin and Garraffo 2005; Byers and Pringle 2006; Robinson and Brink 2006). The optimum egg size (and thus pelagic larval duration) with no dispersal is determined by the values of *S*_crit_, *g*, and *m* (eq. [4]). *S*_crit_ and *g* were set to be comparable with Levitan (2000) and Pringle *et al*. (2014), and *m* was adjusted accordingly to give an optimal pelagic larval duration with no dispersal around 40 days so that evolutionary stable pelagic larval durations would be within realistic ranges for benthic marine organisms (Shanks et al. 2003; Shanks 2009). Coastline length was set to 100 kilometers. Parameter choices for the number of microsites *K* and total amount of material contributed to egg production *C* did not affect evolutionary outcomes and were arbitrary set to give reasonable numbers of larvae. The sensitivity of results to parameter values is presented in the results.

## Results

### Evolution of pelagic larval duration (*T*_PLD_)

#### No mean alongshore flow

Without ocean currents (*i.e.*, *U*=0 and *σ*=0), there is only one pelagic larval duration (*T*_PLD_) that is an evolutionarily stable strategy (ESS) and it occurs at the *T*_PLD_ which maximizes the total number of larvae that survive the pelagic phase (Figure 1). This result recovers classic results from previous models without oceanography (Smith and Fretwell 1974; Levitan 2000).

If there are short timescale fluctuations in flow, but still no mean alongshore flow (*i.e. U*=0 and *σ*>0), the ESS *T*_PLD_ will slightly decrease from the optimum shown in figure 1A (Figure 2 when scaled alongshore flow is 0). The slight decrease occurs because our model assumes there is a limited range of habitable coastline and a higher *T*_PLD_ leads to more individuals dispersing away from the source location and thus being lost off the upstream or downstream edge of the habitable range. However, this effect is small. Given our baseline parameter values, the ESS *T*_PLD_ never decreased more than 3 days even given the highest values of *σ* we would expect to see in nature.

**Figure 2.**
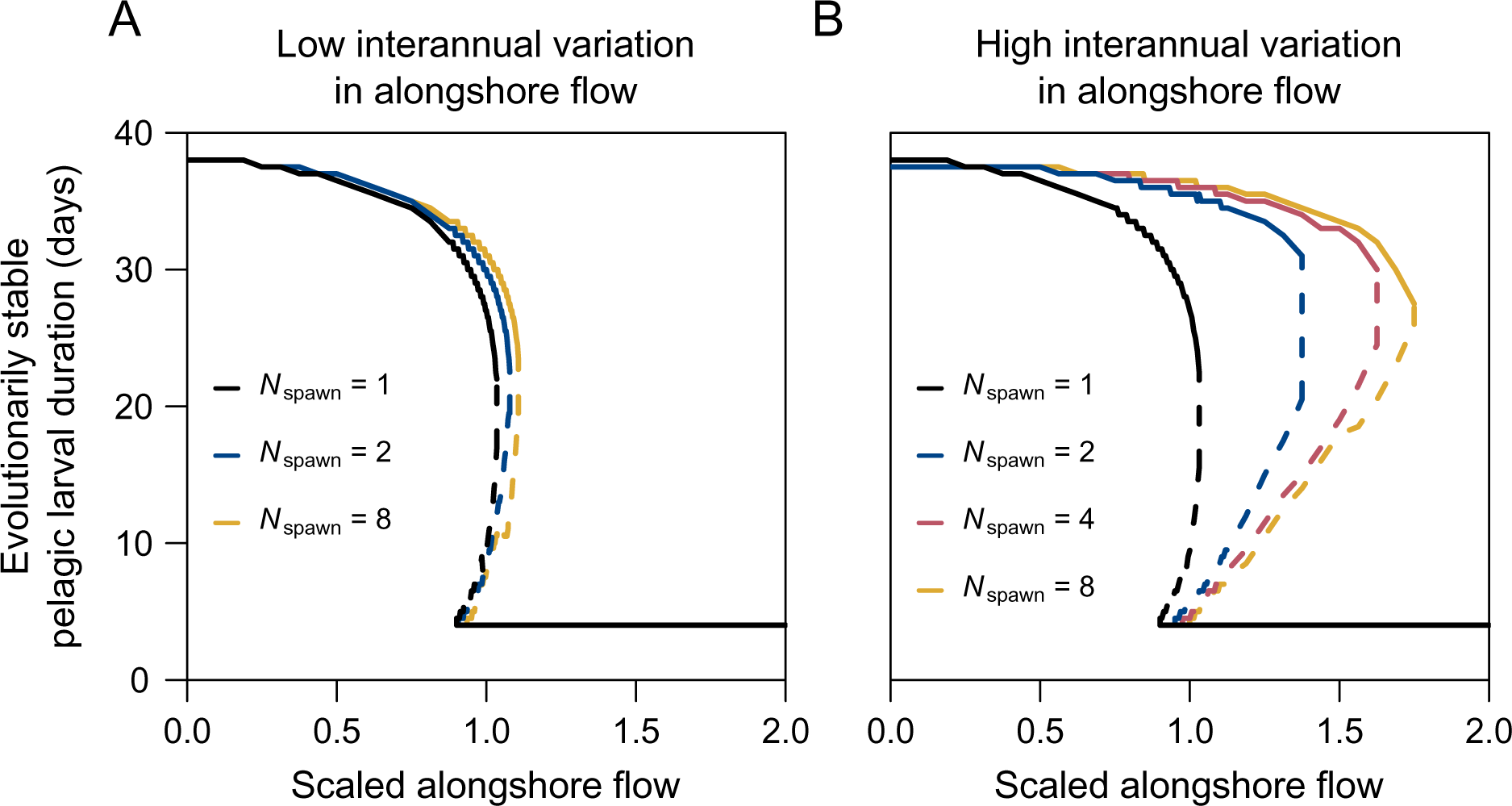
Effect of scaled alongshore flow and the number of spawning events (*N*_spawn_) on the evolutionary stable pelagic larval duration (*T*_PLD_). Panel A shows results when the standard deviation of interannual mean flow rates is *σ*_IA_=0.012 (meters/second) and panel B shows results for when *σ*_IA_=0.035 (meters/second). Solid lines denote evolutionary stable values of *T*_PLD_ while dashed lines denote unstable equilibrium values of *T*_PLD_. Different colors denote different numbers of spawning events (*N*_spawn_) as indicated in the figure legend. Parameters: *σ*=8000 (meters/day), *τ*=4, C=0.1, *s*_crit_=0.0103, *g* = 0.16, *m* =5×10^-7^, *A_m_*=0.1, *K*=200, length of coastline = 100 kilometers, minimum value of *T*_PLD_ was set to *τ*.

#### Effects of scaled alongshore flow on the evolution of pelagic larval duration

As the mean alongshore flow (*U*) current increases, relative to short timescale fluctuations in currents (*σ*), which we refer to as “scaled alongshore flow”, there is selection for decreased *T*_PLD_ (Figure 2). For small to moderate values of scaled alongshore flow (⪅1), there is only one evolutionary stable *T*_PLD_, the value of which is relatively high but decreases as scaled alongshore flow increases for a given number of spawning events. For high values of scaled alongshore flow (⪆1), there are two evolutionary stable *T*_PLD_ values, one of which is at the minimum possible *T*_PLD_ (we set this minimum *T*_PLD_ = *τ*). The two ESSs are separated by an unstable equilibrium (indicated by the dashed lines in Figure 2 for each *N*_spawn_). If the population with a given *N*_spawn_ begins with a *T*_PLD_ below this unstable equilibrium, evolution by a series of small mutations will lead the population to the lower evolutionary stable *T*_PLD_. Alternatively, if the population begins with a *T*_PLD_ above this unstable equilibrium, evolution by a series of small mutations will lead the population to the higher evolutionary stable *T*_PLD_. Figure 3 shows pairwise invasibility plots for examples when there is one ESS or two ESSs for *T*_PLD_.

**Figure 3.**
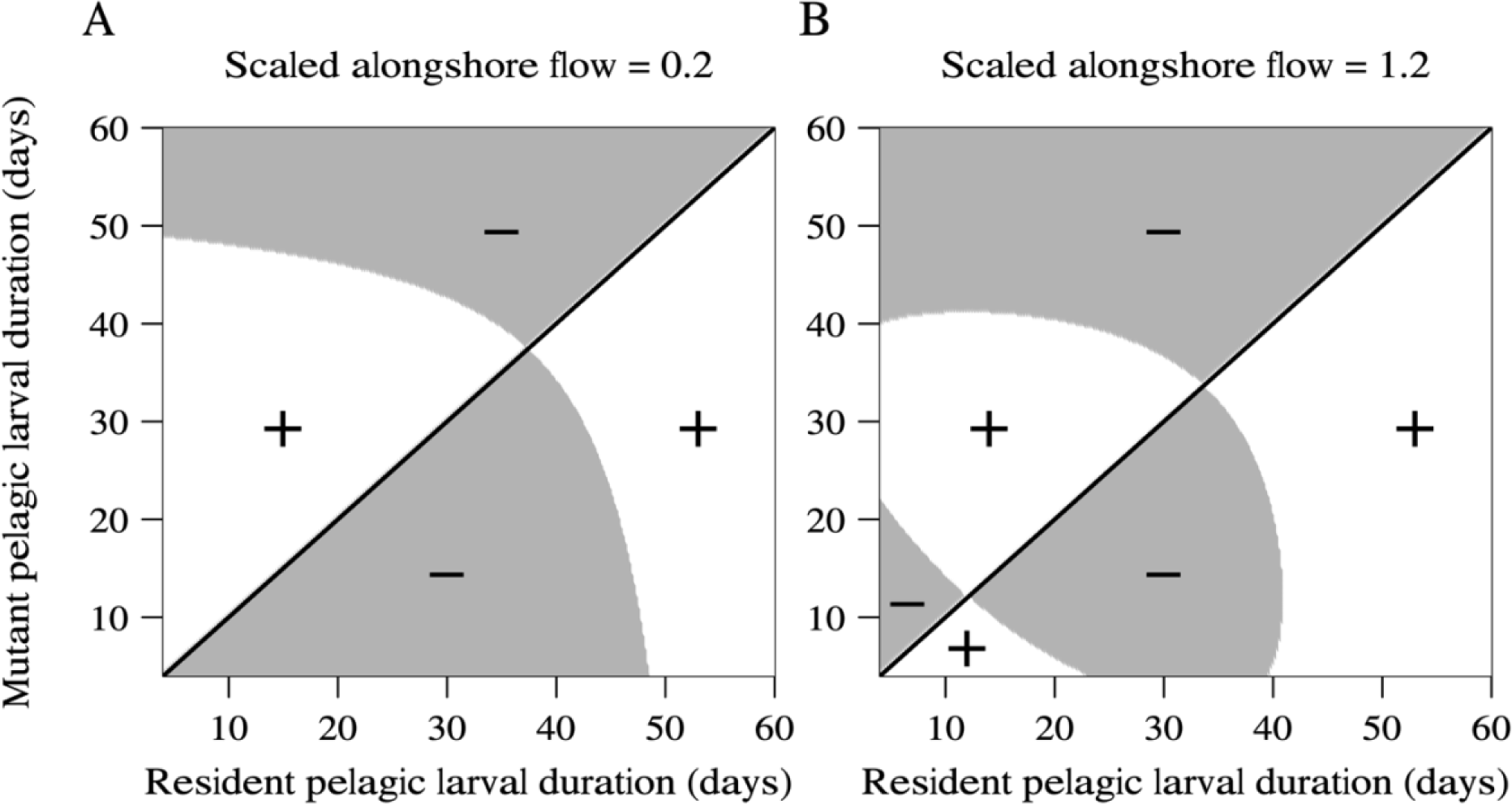
Example pairwise invasibility plots for when there is one ESS (left panel) or two ESSs (right panel). Invasion fitness of the mutant is greater than 1 in the white areas and less than 1 in the grey areas. The black line denotes the 1:1 line where the invasion fitness of the mutant is 1.0. In both plots, *N*_spawn_ = 2 and all parameters are the same as in figure 2B.

These general results presented as evolutionary stable strategies for semelparous organisms are qualitatively similar to what Pringle et al (2014) showed in terms of selection coefficients in non-overlapping generations. There is selection for the pelagic larval durations that lead to the most larvae at or upstream of the source location (Pringle et al. 2014). With low scaled alongshore flow, enough local retention occurs with both short and long *T*_PLD_, but a longer *T*_PLD_ maximizes fecundity because it is associated with the production of smaller, more numerous eggs. In contrast, with high scaled alongshore flow, a long *T*_PLD_ results in insufficient local retention to offset the losses of larvae downstream. As a result, upstream retention is greatest with a short *T*_PLD_ because it minimizes downstream dispersal. However, unlike Pringle et al. (2014) we show a continuous decline in ESS *T*_PLD_ as scaled alongshore flow increases up until the ESS *T*_PLD_ reaches the minimum value. In other words, for any given scaled alongshore flow below a certain strength (∼1.0 in with our parameters), there is a unique ESS *T*_PLD_ (Figure 2). This differs from the results of Pringle et al. (2014) which only predicted two possible evolutionary stable pelagic larval durations: the maximum possible value and the minimum possible value.

#### Effects of the number of spawning events on the evolution of pelagic larval duration

The number of spawning events (*N*_spawn_) changes the pelagic larval duration that is evolutionarily stable under a given flow regime. Spawning more often allows longer pelagic larval durations to remain evolutionarily stable for higher mean scaled alongshore flows (Figure 2). For instance, consider a mean scaled alongshore flow of 1.5 in Figure 2B. If *N*_spawn_*=*1, the only ESS *T*_PLD_ is at 4 days (the minimum *T*_PLD_), but if *N*_spawn_*=*8, a *T*_PLD_ of 33.5 days is evolutionarily stable as well as *T*_PLD_ of 4 days. As interannual variation in mean flow rates increases (*σ*_IA_ > 0), increasing *N*_spawn_ allows for higher ESS values of *T*_PLD_. Increasing *N*_spawn_ increases the range of scaled alongshore flow rates where there are two ESS *T*_PLD_ values because spawning on multiple occasions has a smaller effect on the lower evolutionarily stable pelagic larval duration than then upper one (Figure 2).

The reasons why longer pelagic larval durations become evolutionarily stable when spawning is more frequent can be understood in terms of what Byers and Pringle (2006) showed for population persistence. Larvae released in different years are exposed to different mean alongshore current speeds and directions. Parents that release larvae in multiple years have greater dispersion in their lifetime dispersal kernels (the dispersal kernel of all larvae released over an individual’s lifespan). Increased dispersion allows parents to retain more larvae at or upstream of their spawning location without necessarily increasing fecundity. However, longer pelagic larval durations both increase dispersion and increase fecundity. Therefore, in our adaptive dynamics framework, spawning more frequently (increasing dispersion and upstream retention), together with the effect of longer pelagic larval durations (increasing dispersion and fecundity), allows longer *T*_PLD_ to become evolutionarily stable in addition to short pelagic larval durations. In contrast, short pelagic larval durations are the only evolutionarily stable strategy in non-overlapping generations with high scaled alongshore flow.

### Evolution of the number of spawning events (*N*_spawn_)

Larger interannual fluctuations typically lead to greater evolutionarily stable numbers of spawning events (Figure 4, but see Figure A2B for a counter example). In addition, longer pelagic larval durations and greater scaled alongshore flow speeds also lead to greater evolutionarily stable numbers of spawning events (Figure 4). At low scaled alongshore flow, the evolutionarily stable number of spawning events is always 1 because increasing *N*_spawn_ increases the diffusion experienced over the lifetime on an individual (increasing *L*_diffeffect_ when *σ*_IA_>0), and increases the number of offspring dispersing away from the source location which reduces local growth rates. As scaled alongshore flow increases however, the evolutionarily stable number of spawning events increases to offset the increased loss of larvae downstream, and it increases quicker with longer pelagic larval durations (Figure 4). Longer pelagic larval durations lead to greater downstream losses, more so as scaled alongshore flow increases. Therefore, more frequent spawning evolves to increase retention and offset the downstream losses from increasingly longer pelagic larval durations.

**Figure 4.**
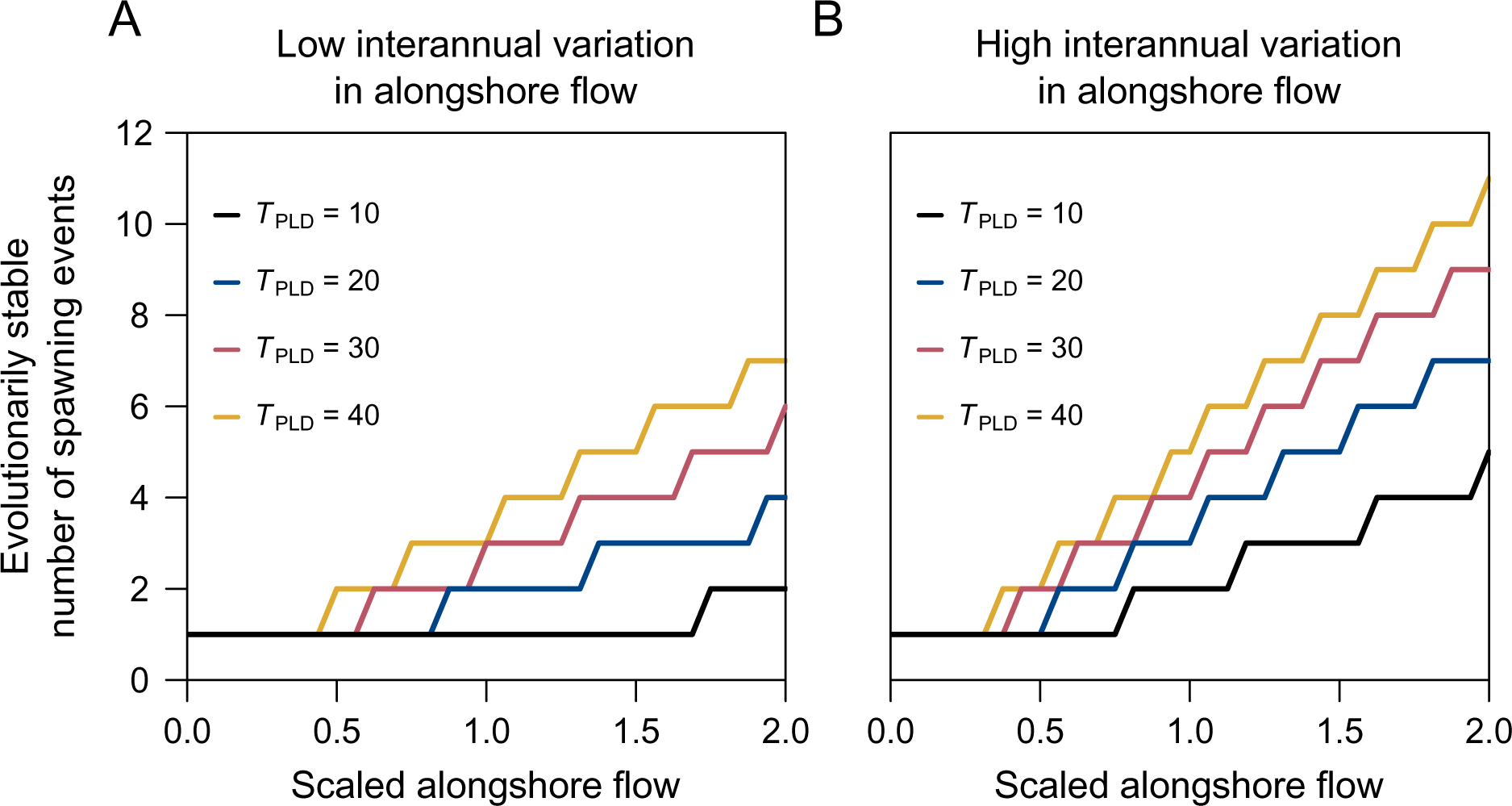
Effect of scaled alongshore flow and pelagic larval duration (*T*_PLD_) on the evolutionarily stable number of spawning events (*N*_spawn_) when there is adult mortality *A_m_*=0.1. Panel A shows results when the standard deviation of interannual mean flow rates *σ*_IA_=0.012 meters/second and panel B shows results for when *σ*_IA_=0.035 meters/second. Different colors denote different pelagic larval durations as indicated in the figure legend. All other parameters are the same as in figure 2.

### Coevolution of pelagic larval duration and the number of spawning events

When pelagic larval duration and the number of spawning events coevolve, three insights emerge (Figure 5). First, when scaled alongshore flow is low, a unique long pelagic larval duration and a unique small number of spawning events are evolutionarily stable, and are similar to when they evolve independently. Second, when scaled alongshore flow is high, a unique short pelagic larval duration and a unique large number of spawning events are evolutionarily stable, also similar to when they evolve independently. Third, and in contrast to outcomes when each trait evolves independently, at intermediate scaled alongshore flows, two combinations of pelagic larval duration and number of spawning events are evolutionarily stable: one with a long pelagic larval duration and many spawning events, and another with a short pelagic larval duration and few spawning events (Figure 5). In particular, two evolutionarily stable numbers of spawning events at higher scaled alongshore flow were not predicted when considering the evolution of the number of spawning events in isolation (Figure 4). With greater degrees of interannual variation, there is a greater range of scaled alongshore flow values that lead to two evolutionarily stable combinations of pelagic larval duration and number of spawning events. The initial trait values determine whether the traits evolve to the upper of lower ESSs (Figure A4)

**Figure 5.**
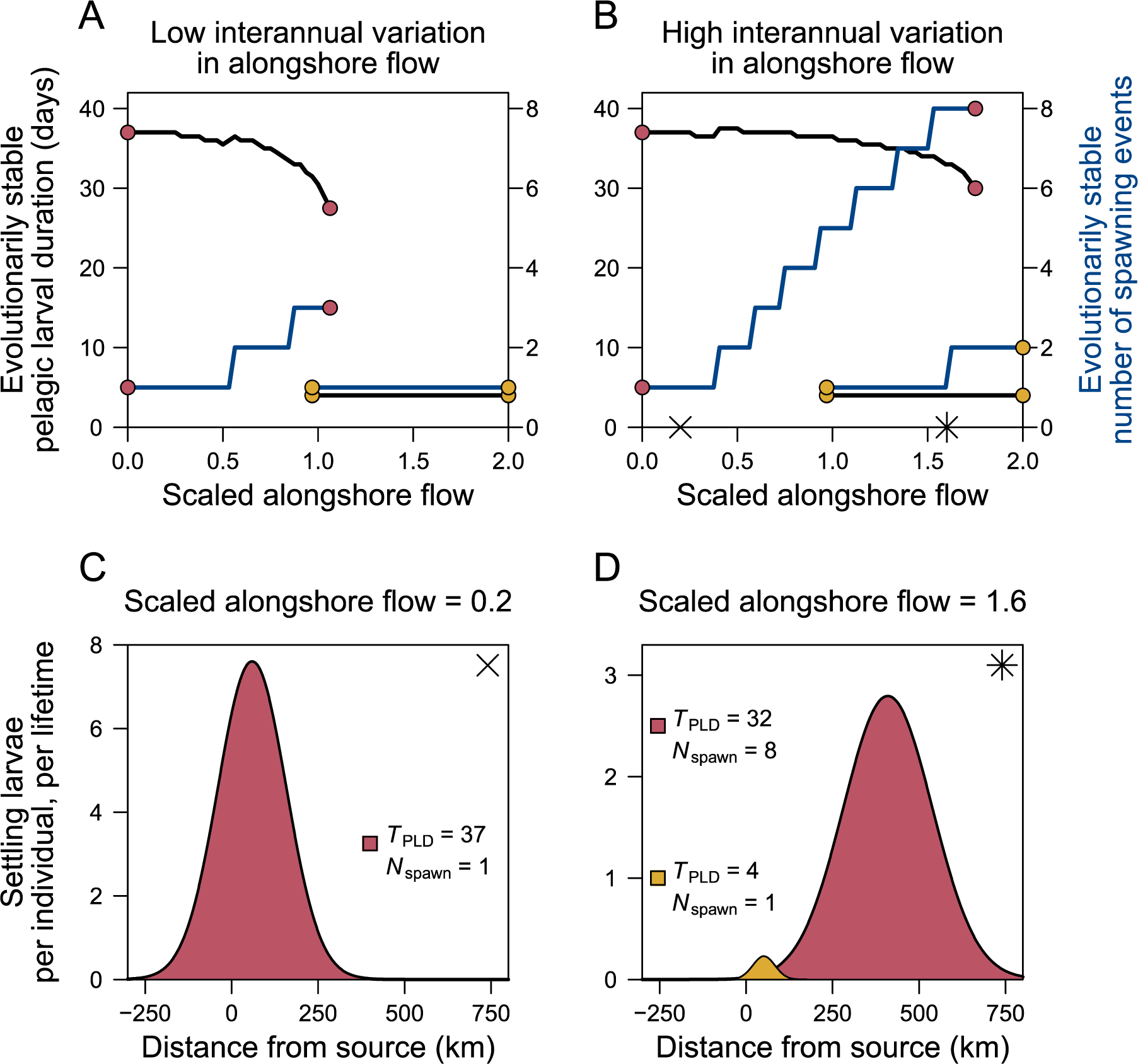
Coevolution of pelagic larval duration (*T*_PLD_) and number of spawning events (*N*_spawn_). Top panels show the evolutionarily stable values of *T*_PLD_ (black lines) and *N*_spawn_ (blue lines) for different degrees of scaled alongshore flow with a low standard deviation of interannual mean flow rates (A; *σ*_IA_=0.012 meters/second) or high standard deviation of interannual mean flow rates (B; *σ*_IA_=0.035 meters/second). Lines denoted with the same color circles are evolutionarily stable together. Note that at intermediate values of scaled alongshore flow there are two combinations of *T*_PLD_ and *N*_spawn_ that are evolutionarily stable: one where both *T*_PLD_ and *N*_spawn_ are at high values and where they are both at low values. Bottom panels show the distribution of larvae released by one individual over its lifetime for different evolutionarily stable life history strategies that emerge with high interannual variation in alongshore flow. Panel C shows a case with low scaled alongshore flow when there is only one evolutionarily stable life history and panel D shows a case with intermediate scaled alongshore flow when there are two evolutionarily stable life history strategies. Note difference in scale bars between panels C and D. Parameters the same as in Figure 2.

An outcome of coevolution at low scaled alongshore flow values is that the evolutionarily stable pelagic larval duration does not always decrease monotonically with increased scaled alongshore flow as it does when pelagic larval duration evolves independently (compare Figure 2 to Figure 5A–B). The ESS *T*_PLD_ can increase with an increased scaled alongshore flow (e.g., scaled advection ∼0.5–0.7 in Figure 5A) when selection favors in increase in spawning frequency, which then allows a higher pelagic larval duration to be stable in that current (Figure 4). Likewise, when selection for decreased pelagic larval duration with increased scaled alongshore flow outweighs selection for increased spawning frequency, the ESS *N*_spawn_ can decrease with increased flow rates (Figure 5A–B).

At intermediate values of scaled alongshore flow where two combinations of pelagic larval duration and number of spawning events are evolutionarily stable, the evolution of spawning events effects the maximum value of scaled alongshore flow where the upper ESS combination is stable. As seen in Figure 2, if only *T*_PLD_ evolves, the upper ESS remains stable with higher scaled alongshore flow if there is a higher value of *N*_spawn_. Therefore, if there is a higher ESS *N*_spawn_ for a given scaled alongshore flow in the coevolutionary model, the upper ESS *N*_spawn_ values remain stable with more scaled alongshore flow. In contrast, the evolution of spawning frequency has little effect on pelagic larval duration in the lower ESS combination. The lower ESS *T*_PLD_ is always the minimum possible value and thus the lower ESS *N*_spawn_ evolves how it does when it evolved independently with a *T*_PLD_ fixed at this value (Figure 4). At the highest values of scaled alongshore flow, the upper ESS combination is no longer stable and only the lower combination of traits remains.

#### Sensitivity to parameter values

The qualitative effects of changes in scaled alongshore flow on life history and dispersal evolution are robust to changes of parameter values in our model. Increasing adult mortality rate (*A_m_*) hampers selection to increase spawning frequency (Figure A5) because higher adult mortality rate discounts the benefits of successive spawning in terms of increasing the standard deviation of dispersal distance, and also results in reduced lifetime fecundity because individuals might die between spawning events. As a result, adult mortality can lead to lower ESS pelagic larval durations in situations where a higher ESS *T*_PLD_ value is only evolutionarily stable when there is a high spawning frequency (Figure A5). Moreover, increasing *A_m_* also selects for shorter pelagic larval durations by reducing recruitment competition (because fewer individuals survive to reproduce) and thus decreasing the benefit of high fecundity, as well as by decreasing the width of the lifetime dispersal kernel. Taken together, the effects of adult mortality can lead to seemingly counterintuitive patterns of *T*_PLD_ evolution. For example, in Figure A5B, the ESS *T*_PLD_ decreases dramatically as scaled alongshore flow increases from low to intermediate rates, because high adult mortality selects against increasing *N*_spawn_ which would allow for higher ESS *T*_PLD_ values without adult mortality. However, as scaled alongshore flow increases further, selection for upstream retention becomes strong enough that the benefits of a larger lifetime dispersal kernel that come with increased *N*_spawn_ outweigh the fecundity cost. Therefore, there is a higher ESS *N*_spawn_, which allows the ESS *T*_PLD_ to dramatically increase (Figure A5B). Finally, with very high scaled alongshore flow, the higher ESS *T*_PLD_ can no longer be maintained even with a high *N*_spawn_ value and thus ESS *T*_PLD_ once again rapidly declines.

Increasing larval growth rate (*g*) or mortality rate (*m)* lowers the ESS *T*_PLD_ and shifts the range of scaled alongshore flow that leads to two evolutionarily stable life history strategies to higher values (Figure A6 and A7). Both of these effects occur because increasing either larval growth rate or mortality rate decreases the *T*_PLD_ that maximizes fecundity (eq. [4]). Changing coastline length had little effect on the coevolutionary outcomes of our model (Figure A8) and changing the number of available subsites *K* or total parental investment in offspring *C* had no effect on our results.

#### Effects of coevolution of pelagic larval duration and the number of spawning events on lifetime dispersal kernels

The theory presented here shows how three key aspects of coastal oceanographic regimes (mean alongshore currents *U*, short-term stochasticity in currents *σ*, and interannual variation in mean alongshore currents *σ*_IA_) select on both larval and adult traits to increase upstream retention and alter the expected mean and spread of lifetime dispersal kernels.

When scaled alongshore flow is low so the potential for movement is low, organisms evolve a long pelagic larval duration and a single spawning release, which results in a high fecundity and dispersal kernels with a mean shifted downstream and high variance (e.g., Figure 5C). When scaled alongshore flow is high so the potential for movement is high, organisms evolve short pelagic larval durations and fewer spawning events, which results in a low fecundity and evolutionarily stable dispersal kernels with a low downstream mean and low variance (*e.g.*, orange distribution in Figure 5D). However, when interannual variation in mean alongshore currents is high, a second dispersal kernel with a mean distance shifted downstream and a much higher spread in dispersal distances is also evolutionarily stable (*e.g.*, red distribution in Figure 5D). This second dispersal kernel is the result of selection for a long pelagic larval duration (which selects for small eggs sizes and thus high fecundity, eq. [2]) and high spawning frequency, which is just another way to counter the downstream losses of larvae, but results in much greater mean and variance in dispersal distances as a consequence.

## Discussion

We sought to understand how coastal oceanographic processes affect the evolution of marine life history traits and how that could indirectly affect the expected distribution of marine dispersal distances. Along most coastlines, there is usually a dominant water flow direction that biases larval dispersal downstream, and stochastic events during dispersal (like eddies and weather), as well as seasonal and yearly changes in mean flow speed and direction, that slow or reverse currents allowing occasional upstream retention (Largier 2003; Lumpkin and Garraffo 2005; Shanks and Eckert 2005). These common features of coastal environments act as agents of selection on marine life history traits that affect dispersal and larval development, and could potentially explain the evolution of dispersal without invoking the traditional causes of inbreeding, kin competition, and environmental variability. The new and key results emerging from our theory are especially relevant on coastlines with relatively high mean flow rates and high interannual variation in flow rates (Largier 2003; Lumpkin and Garraffo 2005). First, selection induced by coastal oceanography favors the release of larvae over multiple time periods, rather than all at once. Releasing larvae on multiple occasions allows individuals to avoid extinction from net downstream larval loss by increasing the variance in their lifetime dispersal kernel. Doing so reduces the fitness costs of long pelagic larval durations predicted in Pringle et al (2013). Costs are reduced by offsetting downstream losses under strong currents, allowing long pelagic larval durations to be maintained in marine life cycles if it allows individuals to access greater fecundity through reduced parental investment per offspring. Second, while pelagic larval duration and the number of spawning events both affect dispersal, the evolution of the number of spawning events affects the evolution of pelagic larval duration, and vice versa. Such coevolution between larval and adult traits changes how currents affect the evolution of each trait separately and the expected dispersal distances that evolve in a given current regime. Third, the same current regime can give rise to populations with quite different evolutionarily stable pelagic larval durations and spawning frequencies. Finally, the evolution of quite different pelagic larval durations and spawning frequency gives rise to dispersal kernels with very different means and variances in dispersal distances. Our model is structured in such a way that it can be parameterized with data to explore specific situations. The main implication of our findings is that coastal ocean flows are important agents of selection that can generate multiple, often co-occurring, evolutionary outcomes for marine life history traits that affect dispersal.

Our findings offer a new explanation for the disconnect between the diversity of pelagic larval durations and spawning frequencies found co-occurring in nature and the predictions from classic marine life history theory that species should produce either many small eggs or few large eggs (Vance 1973), or a single intermediate optimal egg sizes depending on larval growth and mortality rates (e.g., Levitan 2000). A given combination of larval growth and mortality rates can lead to a range of evolutionarily stable life history strategies depending on the oceanographic conditions during the time in which larvae are released. Our adaptive dynamics approach allows us to identify oceanographic conditions where two different life history strategies are evolutionarily stable, and which one evolves depends on a population’s evolutionary starting point. For instance, closely related species might evolve very different pelagic larval durations (and thus different egg sizes) and spawning frequencies if they live on coastlines with different currents, or spawn at different times of the year with different currents. Moreover, even on the same section of coast, similar species could evolve dramatically different pelagic larval durations and spawning frequencies simply because of their different evolutionary histories (although the coexistence of these two species would require niche differences if there are no fitness differences). Combining these effects with among-species differences in larval growth rates, larval mortality rates, adult mortality rates, and other parameters, all of which lead to different evolutionarily stable life histories, it becomes clearer how a diversity of life histories can be seen in nature on any given stretch of coastline.

By showing how dispersal kernels can be shaped by the coevolution of larval and adult traits, our results imply that considering either larval or adult traits in isolation might produce incorrect predictions about how life history traits and dispersal kernels evolve. Previous marine dispersal theory has either modeled the dispersal kernel inherently as an unconstrained trait responding to habitat heterogeneity (Shaw et al. 2019) or only modeled selection on larval traits (Pringle et al. 2014). Byers and Pringle (2006) showed that spawning over multiple time periods could increase population persistence and spread through upstream retention but did not consider the evolution of either pelagic larval duration or spawning frequency. Shanks and Eckert (2005) compiled data on nearshore and shelf/slope fishes and crustaceans and made the case that both adult traits (e.g., longevity, the number of broods per year) and larval traits (e.g., pelagic larval durations) have evolved to exploit eddies and counter-currents to aid in larval retention. Our theory provides a framework to understand how selection for larval retention influences the evolution of both larval and adult traits.

In the presented form, our model makes qualitive predictions about the evolution of marine life histories for a board range of realistic parameters, but its integral projection model structure makes it easily adaptable to match specific systems. When populations are structured in multiple dimensions (e.g., space and age), integral projection models typically require the estimation of many fewer parameters than an equivalent matrix population model (Ellner and Rees 2006). Empirical estimates of the parameters in our model could be used to give specific predictions about evolutionary outcomes in specific situations. Perhaps more usefully, however, any of the functions that give transition probabilities between stages could also be replaced with empirically estimated relationships. For instance, we assume a specific relationship between egg size and the probability of surviving the pelagic larval stage (eq. 1–3). Researchers interested in a specific species could instead estimate this relationship by collecting data and fitting a statistical model, such as (Graham et al. 2008; Connolly and Baird 2010; Moneghetti et al. 2019). This new estimated function could then replace equation 3 and thus *f* in equation 8. However, it is important to note, that when applying our model to real systems, researchers should take care to estimate oceanographic statistics on a spatiotemporal scale relevant to their study species (Largier 2003). For instance, the annual mean alongshore flow rate might be an inappropriate measure for predicting life history evolution of a species that only spawns in April each year.

Many of the qualitative predictions from our model match empirical patterns. For instance, the prediction that there should be shorter pelagic larval durations with stronger scaled alongshore flow is supported by evidence that the proportion of marine invertebrate species with planktotrophic larvae decreases with scaled alongshore flow rate (Marshall et al. 2012; Pringle et al. 2014; Álvarez-Noriega et al. 2020). Our model’s prediction that, for many different oceanographic conditions, there are two different evolutionary stable pelagic larval durations (one long and one short) is consistent with previous data compilations that pelagic larval durations are bimodally distributed among benthic marine species, with dispersal distances <1KM or >∼20km (Shanks et al. 2003; Shanks 2009). While the dispersal distances <1km in these datasets include short-lived, non-feeding larvae, they also include longer-lived feeding larvae, and our model predicts that bimodality can emerge even if all species have feeding larvae. Data from the fishes and crustaceans off the coast of California also support our predictions that species will have longer pelagic larval durations with greater short-time scale fluctuations in alongshore flow and higher spawning frequencies with greater inter-spawning-event variation in alongshore flow (Shanks and Eckert 2005). Shanks and Eckert (2005) also found a positive correlation between maximum age and pelagic larval duration, which matches our coevolutionary predictions if living longer equates to more spawning events. Empirical studies also emphasize a factor not included in our model, the timing of spawning (e.g., Morgan and Christy 1995; Reitzel et al. 2004; Shanks and Eckert 2005), which affects the scaled alongshore flow and interannual variation experienced, finding that a disproportionate number of species have evolved to spawn during seasons with relatively low alongshore flow rates or across months when currents reverse directions (Shanks and Eckert 2005; Byers and Pringle 2006). Other factors not included in our model are cross-shore currents and larval swimming behavior which could interact to affect the realized scaled alongshore flow (Largier 2003; Meyer et al. 2021*a*; Morgan et al. 2021). In addition, our model assumes that pelagic larval duration and the number of spawning events are genetically independent traits. While there is some evidence for genetic correlations in larval and adults traits in marine organisms (e.g., Levin et al. 1991; Miles and Wayne 2009; Ciosi et al. 2011), and robust theory for how such correlations are expected to influence evolutionary trajectories and rate of adaptation (e.g., Marshall and Connallon 2023), their long term effect on the outcome of evolution, which our model focuses on, remains less clear (Gomulkiewicz and Houle 2009; Hansen et al. 2023). However, like all simplifications of complex phenomena, our model serves the purpose of re-orientating and focusing empirical research, and learning why observations match or do not match model predictions. In particular, it provides predictions of the parameter space where larval behaviors would have greater or less impact and how they could possibly substitute for the role of pelagic larval duration or spawning frequency.

In the future, our model could be extended to include other concepts from the marine dispersal literature such as non-Gaussian dispersal kernels (Pringle et al. 2009; Chiswell 2012; Stover et al. 2014), non-feeding larvae (Marshall and Bolton 2007; Marshall and Keough 2007), swimming behavior (Meyer et al. 2021*a*; Burgess et al. 2022), or spatial heterogeneity in habitat availability or quality (Baskett et al. 2007; Meyer et al. 2021*b*). We expect that incorporating the evolution of additional traits such as timing of spawning or larval swimming behavior into our model would allow longer pelagic larval durations to be evolutionarily stable on coastlines with strong directional currents because such traits typically function to increase upstream retention (Paris and Cowen 2004; Bottesch et al. 2016; Morgan et al. 2021; Burgess et al. 2022). Existing dispersal theory predicts that increased spatial heterogeneity, kin selection, and inbreeding depression select for increased dispersal (Clobert et al. 2012), however, researchers have not yet evaluated how these factors will interact with the life history trade-offs and oceanographic effects that are central to our results. Ultimately, a comprehensive theory of dispersal evolution, applicable to both terrestrial and marine organisms, will integrate the ideas discussed here that focus on the evolution of traits that give rise to dispersal outcomes with key factors in dispersal evolution theory that directly cause selection on dispersal outcomes (e.g., variation in local conditions, kin selection, and inbreeding depression; Clobert et al. 2012).

## Acknowledgements

We are grateful to J. M. Pringle and other members of the Burgess lab who provided helpful feedback on this project. We also thank the editors and anonymous reviewers, whose comments greatly improved the manuscript.

## Literature cited

Álvarez-Noriega, M., S. C. Burgess, J. E. Byers, J. M. Pringle, J. P. Wares, and D. J. Marshall. 2020. Global biogeography of marine dispersal potential. Nature Ecology & Evolution 4:1196–1203.

Baskett, M. L., J. S. Weitz, and S. A. Levin. 2007. The evolution of dispersal in reserve networks. The American Naturalist 170:59–78.

Bottesch, M., G. Gerlach, M. Halbach, A. Bally, M. J. Kingsford, and H. Mouritsen. 2016. A magnetic compass that might help coral reef fish larvae return to their natal reef. Current Biology 26:R1266–R1267.

Brännström, Å., J. Johansson, and N. Von Festenberg. 2013. The hitchhiker’s guide to adaptive dynamics. Games 4:304–328.

Burgess, S. C., M. L. Baskett, R. K. Grosberg, S. G. Morgan, and R. R. Strathmann. 2016. When is dispersal for dispersal? Unifying marine and terrestrial perspectives. Biological Reviews 91:867–882.

Burgess, S. C., M. Bode, J. M. Leis, and L. B. Mason. 2022. Individual variation in marine larval-fish swimming speed and the emergence of dispersal kernels. Oikos 2022:e08896.

Byers, J., and J. Pringle. 2006. Going against the flow: retention, range limits and invasions in advective environments. Marine Ecology Progress Series 313:27–41.

Cecino, G., and E. A. Treml. 2021. Local connections and the larval competency strongly influence marine metapopulation persistence. Ecological Applications 31:e02302.

Chesson, P. L., and R. R. Warner. 1981. Environmental Variability Promotes Coexistence in Lottery Competitive Systems. The American Naturalist 117:923–943.

Chiswell, S. M. 2012. Non-Gaussian larval dispersal kernels in Gaussian ocean flows. Aquatic Biology 16:203–208.

Ciosi, M., N. J. Miller, S. Toepfer, A. Estoup, and T. Guillemaud. 2011. Stratified dispersal and increasing genetic variation during the invasion of central europe by the western corn *rootworm, diabrotica virgifera virgifera*. Evolutionary Applications 4:54–70.

Clobert, J., M. Baguette, T. G. Benton, and J. M. Bullock, eds. 2012. Dispersal Ecology and Evolution. Oxford University Press, Oxford, United Kingdom.

Connolly, S. R., and A. H. Baird. 2010. Estimating dispersal potential for marine larvae: dynamic models applied to scleractinian corals. Ecology 91:3572–3583.

Davis, R. E. 1985. Drifter observations of coastal surface currents during CODE: The statistical and dynamical views. Journal of Geophysical Research: Oceans 90:4756–4772.

Dewi, S., and P. Chesson. 2003. The age-structured lottery model. Theoretical Population Biology, Understanding the role of environmental vatiation in population and community dynamics 64:331–343.

Ellner, S. P., and M. Rees. 2006. Integral projection models for species with complex demography. The American Naturalist 167:410–428.

Emlet, R. B., L. R. McEdward, and R. R. Strathmann. 1987. Echinoderm larval ecology viewed from the egg. Pages 55–136 in M. Jangoux and J. M. Lawrence, eds. Echinoderm studies (Vol. 2). CRC Press, London, UK.

Gaylord, B., and S. D. Gaines. 2000. Temperature or Transport? Range Limits in Marine Species Mediated Solely by Flow. The American Naturalist 155:769–789.

Gomulkiewicz, R., and D. Houle. 2009. Demographic and Genetic Constraints on Evolution. The American Naturalist 174:E218–E229.

Graham, E. M., A. H. Baird, and S. R. Connolly. 2008. Survival dynamics of scleractinian coral larvae and implications for dispersal. Coral Reefs 27:529–539.

Grantham, B. A., G. L. Eckert, and A. L. Shanks. 2003. Dispersal potential of marine invertebrates in diverse habitats. Ecological Applications 13:108–116.

Hansen, T. F., D. Houle, M. Pavličev, and C. Pélabon, eds. 2023. Evolvability: A Unifying Concept in Evolutionary Biology? MIT Press, Cambridge, Massachusetts, USA.

Iwasa, Y., Y. Yusa, and S. Yamaguchi. 2022. Evolutionary game of life-cycle types in marine benthic invertebrates: Feeding larvae versus nonfeeding larvae versus direct development. Journal of Theoretical Biology 537:111019.

Jackson, G. A., and R. R. Strathmann. 1981. Larval mortality from offshore mixing as a link between precompetent and competent periods of development. The American Naturalist 118:16–26.

Kinlan, B. P., and S. D. Gaines. 2003. Propagule dispersal in marine and terrestrial environments: a community perspective. Ecology 84:2007–2020.

Largier, J. L. 2003. Considerations in estimating larval dispersal distances from oceanographic data. Ecological Applications 13:71–89.

Leis, J. M. 2006. Are larvae of demersal fishes plankton or nekton? Pages 57–141 inAdvances in Marine Biology (Vol. 51). Academic Press.

Levin, L. A., J. Zhu, and E. Creed. 1991. The genetic basis of life-history characters in a polychaete exhibiting planktotrophy and lecithotrophy. Evolution 45:380–397.

Levitan, D. R. 2000. Optimal egg size in marine invertebrates: theory and phylogenetic analysis of the critical relationship between egg size and development time in echinoids. The American Naturalist 156:175–192.

Lumpkin, R., and Z. Garraffo. 2005. Evaluating the decomposition of tropical atlantic drifter observations. Journal of Atmospheric and Oceanic Technology 22:1403–1415.

Marshall, D. J., D. R. Barneche, and C. R. White. 2022. How does spawning frequency scale with body size in marine fishes? Fish and Fisheries 23:316–323.

Marshall, D. J., and T. F. Bolton. 2007. Effects of egg size on the development time of non-feeding larvae. The Biological Bulletin 212:6–11.

Marshall, D. J., and T. Connallon. 2023. Carry-over effects and fitness trade-offs in marine life histories: The costs of complexity for adaptation. Evolutionary Applications 16:474–485.

Marshall, D. J., and M. J. Keough. 2007. The evolutionary ecology of offspring size in marine invertebrates. Advances in Marine Biology 53:1–60.

Marshall, D. J., P. J. Krug, E. K. Kupriyanova, M. Byrne, and R. B. Emlet. 2012. The biogeography of marine invertebrate life histories. Annual Review of Ecology, Evolution, and Systematics 43:97–114.

Marshall, D. J., A. K. Pettersen, and H. Cameron. 2018. A global synthesis of offspring size variation, its eco-evolutionary causes and consequences. Functional Ecology 32:1436– 1446.

McGill, B. J., and J. S. Brown. 2007. Evolutionary game theory and adaptive dynamics of continuous traits. Annual Review of Ecology, Evolution, and Systematics 38:403–435.

Metaxas, A., and M. Saunders. 2009. Quantifying the “bio-” components in biophysical models of larval transport in marine benthic invertebrates: advances and pitfalls. The Biological Bulletin 216:257–272.

Meyer, A. D., A. Hastings, and J. L. Largier. 2021a. Larvae of coastal marine invertebrates enhance their settling success or benefits of planktonic development – but not both – through vertical swimming. Oikos 130:2260–2278.

Meyer, A. D., A. Hastings, and J. L. Largier. 2021b. Spatial heterogeneity of mortality and diffusion rates determines larval delivery to adult habitats for coastal marine populations. Theoretical Ecology 14:525–541.

Miles, C. M., and M. L. Wayne. 2009. Life history trade-offs and response to selection on egg size in the polychaete worm *Hydroides elegans*. Genetica 135:289–298.

Moneghetti, J., J. Figueiredo, A. H. Baird, and S. R. Connolly. 2019. High-frequency sampling and piecewise models reshape dispersal kernels of a common reef coral. Ecology 100:e02730.

Morgan, S. G. 2014. Behaviorally mediated larval transport in upwelling systems. Advances in Oceanography 2014:e364214.

Morgan, S. G., and J. H. Christy. 1995. Adaptive significance of the timing of larval release by crabs. The American Naturalist 145:457–479.

Morgan, S. G., C. D. Dibble, M. G. Susner, T. G. Wolcott, D. L. Wolcott, and J. L. Largier. 2021. Robotic biomimicry demonstrates behavioral control of planktonic dispersal in the sea. Marine Ecology Progress Series 663:51–61.

Müller, K. 1982. The colonization cycle of freshwater insects. Oecologia 52:202–207.

Pachepsky, E., F. Lutscher, R. M. Nisbet, and M. A. Lewis. 2005. Persistence, spread and the drift paradox. Theoretical Population Biology 67:61–73.

Palmer, A. R., and R. R. Strathmann. 1981. Scale of dispersal in varying environments and its implications for life histories of marine invertebrates. Oecologia 48:308–318.

Paris, C. B., and R. K. Cowen. 2004. Direct evidence of a biophysical retention mechanism for coral reef fish larvae. Limnology and Oceanography 49:1964–1979.

Pringle, J., F. Lutscher, and E. Glick. 2009. Going against the flow: effects of non-Gaussian dispersal kernels and reproduction over multiple generations. Marine Ecology Progress Series 377:13–17.

Pringle, J. M., J. E. Byers, P. Pappalardo, J. P. Wares, and D. Marshall. 2014. Circulation constrains the evolution of larval development modes and life histories in the coastal ocean. Ecology 95:1022–1032.

Rees, M., and S. P. Ellner. 2016. Evolving integral projection models: evolutionary demography meets eco-evolutionary dynamics. Methods in Ecology and Evolution 7:157–170.

Reitzel, A. M., B. G. Miner, and L. R. McEdward. 2004. Relationships between spawning date and larval development time for benthic marine invertebrates: a modeling approach. Marine Ecology Progress Series 280:13–23.

Robinson, A. R., and K. H. Brink, eds. 2006. The Sea (Vol. Volume 14A, B. The global coastal ocean: interdisciplinary regional studies and synthesis). Harvard University Press, Cambridge, Massachusetts, USA.

Shanks, A. L. 2009. Pelagic larval duration and dispersal distance revisited. The Biological Bulletin 216:373–385.

Shanks, A. L., and G. L. Eckert. 2005. Population persistence of california current fishes and benthic crustaceans: a marine drift paradox. Ecological Monographs 75:505–524.

Shanks, A. L., B. A. Grantham, and M. H. Carr. 2003. Propagule dispersal distance and the size and spacing of marine reserves. Ecological Applications 13:159–169.

Shaw, A. K., C. C. D’Aloia, and P. M. Buston. 2019. The evolution of marine larval dispersal kernels in spatially structured habitats: analytical models, individual-based simulations, and comparisons with empirical estimates. The American Naturalist 193:424–435.

Siegel, D., B. Kinlan, B. Gaylord, and S. Gaines. 2003. Lagrangian descriptions of marine larval dispersion. Marine Ecology Progress Series 260:83–96.

Smith, C. C., and S. D. Fretwell. 1974. The optimal balance between size and number of offspring. The American Naturalist 108:499–506.

Speirs, D. C., and W. S. C. Gurney. 2001. Population persistence in rivers and estuaries. Ecology 82:1219–1237.

Starrfelt, J., and H. Kokko. 2012. The theory of dispersal under multiple influences. Pages 19–28 inDispersal Ecology and Evolution. Oxford University Press.

Stover, J. P., B. E. Kendall, and R. M. Nisbet. 2014. Consequences of dispersal heterogeneity for population spread and persistence. Bulletin of Mathematical Biology 76:2681–2710.

Strathmann, R. 1974. The spread of sibling larvae of sedentary marine invertebrates. The American Naturalist 108:29–44.

Strathmann, R. 1982. Selection for retention or export of larvae in estuaries. Pages 521–536 in V. S. Kennedy, ed. Estuarine comparisons. Academic Press.

Strathmann, R. R. 1985. Feeding and nonfeeding larval development and life-history evolution in marine invertebrates. Annual Review of Ecology and Systematics 16:339–361.

Strathmann, R. R. 1990. Why life histories evolve differently in the sea. American Zoologist 30:197–207.

Travis, J. M. J., M. Delgado, G. Bocedi, M. Baguette, K. Bartoń, D. Bonte, I. Boulangeat, et al. 2013. Dispersal and species’ responses to climate change. Oikos 122:1532–1540.

Treml, E. A., J. R. Ford, K. P. Black, and S. E. Swearer. 2015. Identifying the key biophysical drivers, connectivity outcomes, and metapopulation consequences of larval dispersal in the sea. Movement Ecology 3:17.

Vance, R. R. 1973. On reproductive strategies in marine benthic invertebrates. The American Naturalist 107:339–352.

Warner, R. R., and P. L. Chesson. 1985. Coexistence Mediated by Recruitment Fluctuations: A Field Guide to the Storage Effect. The American Naturalist 125:769–787.

