## Appendix for "Larval and adult traits coevolve in response to coastal oceanography to shape marine dispersal kernels"

### Numerically evaluating the integral projection model

Following Ellner and Rees (2006), we numerically evaluate our model using the midpoint rule. This method systematically converts the integral projection model into a matrix model by breaking up the continuous functions at mesh points. This model is then evaluated as a spatial- and stage-structured matrix population model (Hunter and Caswell 2005). We define mesh points $x_i$ by dividing the interval 0 to $L$ (the length of the coastline) evenly into $w$ classes and setting $x_i$ to be the midpoint of class $i$:

$$x_i = (1 - 0.5)\,h,$$

where $h = L / w$. The midpoint approximation of equation 8 is then

$$N_0\left(x_j, t+1\right) = r\left(\sum_{a=0}^{N_{\text{spawn}}} h \sum_{i=1}^{w} f(a)\, k\left(x_j, x_i, a\right) N_a\left(x_i, t\right)\right) \qquad \text{(eq. A1).}$$

We can simplify equation 10 by rewriting the second summation as a matrix multiplication such that

$$\mathbf{N}_0(t+1) = r\left(\sum_{a=0}^{N_{\text{spawn}}} \mathbf{K}_a \mathbf{N}_a(t)\right), \qquad \text{(eq. A2)}$$

where $\mathbf{K}_a$ is a matrix whose $(i, j)$ entry is $hf(a)k(x_i, x_j, a)$ and $\mathbf{N}_a(t)$ is a vector whose $i^{th}$ entry is $N_a(x_i, t)$.

As shown by Ellner and Rees (2006), invasion fitness can now be calculated by iterating equation A2 (with the recruitment function given by equation 10) and equation 9. We begin with any nonzero initial distribution of individuals $\mathbf{n}(0)$, which is a matrix giving the number of individuals at each size and age. Let $\mathbf{u}(t) = \mathbf{n}(t) / \|\mathbf{n}(t)\|$, where $\|\mathbf{x}\|$ is the sum of all entries in $\mathbf{x}$. Using equations A2 and 9, we can iterate $\mathbf{u}$(t) until it converges and then $\lambda = \|\mathbf{u}(t)\|$.

**Effect of initial conditions**

The initial values of pelagic larval duration ($T_{PLD}$) and number of spawning events ($N_{spawn}$) determine whether these traits evolve to the upper of lower ESSs (Figure A3). For instance, at intermediate rates of scaled alongshore flow, if the initial $T_{PLD}$ and $N_{spawn}$ values are both low, the population evolves to the lower combination of ESSs. Alternatively, if both traits start at high values, the population evolves to the upper ESSs. In some cases, the relative pace of evolution also matters. For example, consider the case where scaled alongshore flow is intermediate and the initial values of $T_{PLD}$ is high but the initial value of $N_{spawn}$ is low. If $T_{PLD}$ evolves faster than $N_{spawn}$, the population evolves to the lower combination of ESSs (Figure A1, middle column) because this high value of $T_{PLD}$ is only stable for high values of $N_{spawn}$ (Figure 2). However, if both traits evolve at the same rate or evolves $N_{spawn}$ faster, the population evolves to the upper ESSs. This occurs because the high initial value of $T_{PLD}$ selects for a higher $N_{spawn}$ and if $N_{spawn}$ can evolve quick enough the higher ESSs are stable.

**Stochastic simulations**

To check the robustness of our assumption that the resident population is stationary as well as the implications of our estimate of $L_{diffeffect}$ (equation 7), we compared the results of our main model to stochastic simulations which tracked the population dynamics of residents and stochastically varied flow rates. The goal this model was not to fully explore the effects of stochasticity, although that will be a useful goal for future research, rather we aimed to confirm the key qualitative patterns of pelagic larval duration and spawning frequency evolution seen in our deterministic simulations. We did not explore coevolution in these simulations.

Except for the differences mentioned here, the methods for these simulations were the same as those described for the deterministic simulations in the main text. In the deterministic simulations we only tracked the population size of the mutant and assumed that the resident population was fixed at carrying capacity, but in the stochastic simulations we tracked both the resident and mutant population sizes each time step. Additionally, while in the deterministic model we incorporated interannual variation in flow rates by replacing diffusion ($L_{diff}$) with Byers and Pringle's (2006) estimate for the standard deviation of larval dispersal distance for all larvae released over the lifetime of an adult ($L_{\mathrm{diffeffect}}$), for the stochastic simulations mean alongshore flow rate $U$ was drawn from a Gaussian distribution each time step and then used to calculate diffusion $L_{diff}$.

The stochastic simulations confirmed the key qualitative patterns of pelagic larval duration and spawning frequency evolution seen in our deterministic simulations. Figure A9 shows examples of the evolution pelagic larval duration in the stochastic simulations. As with the deterministic simulations, increased scaled advection selected for increased pelagic larval duration and increasing the number of spawning events allowed for a high pelagic larval duration to be evolutionarily stable for higher values of scaled advection (Figure A9). Figure A10 shows examples of the evolution the number of spawning events in the stochastic simulation. As with the deterministic simulations, increasing scaled advection selected for increased spawning frequency, and increasing the pelagic larval duration amplified the effect of scaled advection of the evolution of spawning frequency (Figure A10).

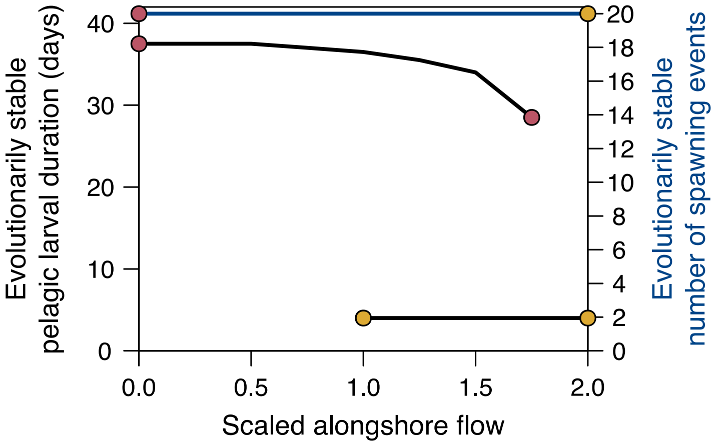


**Figure A1.** Results of coevolutionary model when fecundity is not controlled for by number of spawning events (*i.e.*, $C$ was the same for all spawning events regardless of $N_{spawn}$). The evolutionarily stable values of $T_{\mathrm{PLD}}$ (black lines) and $N_{\mathrm{spawn}}$ (blue lines) for different degrees of scaled alongshore flow with a high standard deviation of interannual mean flow rates ($\sigma_{\mathrm{IA}}$=0.035 meters/second). Populations always evolved to the maximum possible trait value of $N_{\mathrm{spawn}}$, which was set to be 20. Parameters: $\sigma$=8000 (meters/day), $\tau$=4, C=0.1, $s_{\mathrm{crit}}$=0.0103, $g$ = 0.16, $K$=200, $m$=5×10$^{-7}$, $A_m$=0.1, length of coastline = 100 kilometers, minimum value of $T_{\mathrm{PLD}}$ was set to $\tau$.

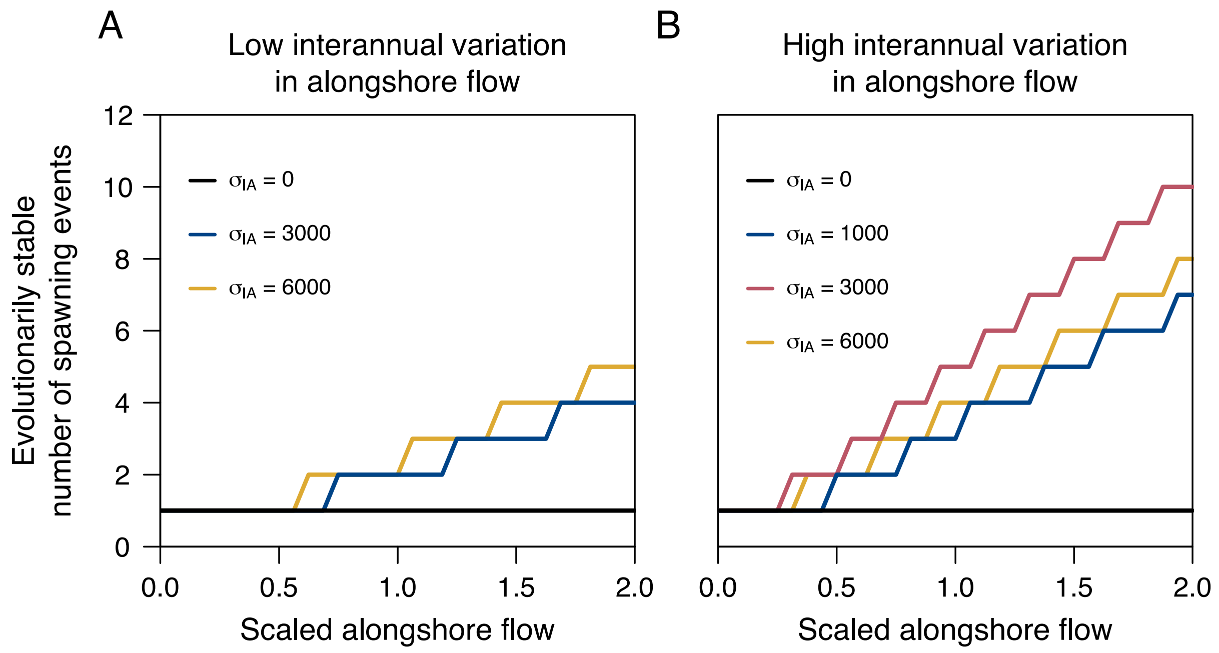


**Figure A2.** Effect of scaled alongshore flow on the evolutionarily stable $N_{spawn}$ when there is no adult mortality $A_m$=0.1. Panel A shows results when the pelagic larval duration is 10 days and panel B shows results for when the pelagic larval duration is 40 days. Different colors denote different standard deviations in interannual mean flow rates $\sigma_{\mathrm{IA}}$ (meters/day) as indicated in the figure legend. All other parameters are the same as in figure 2.

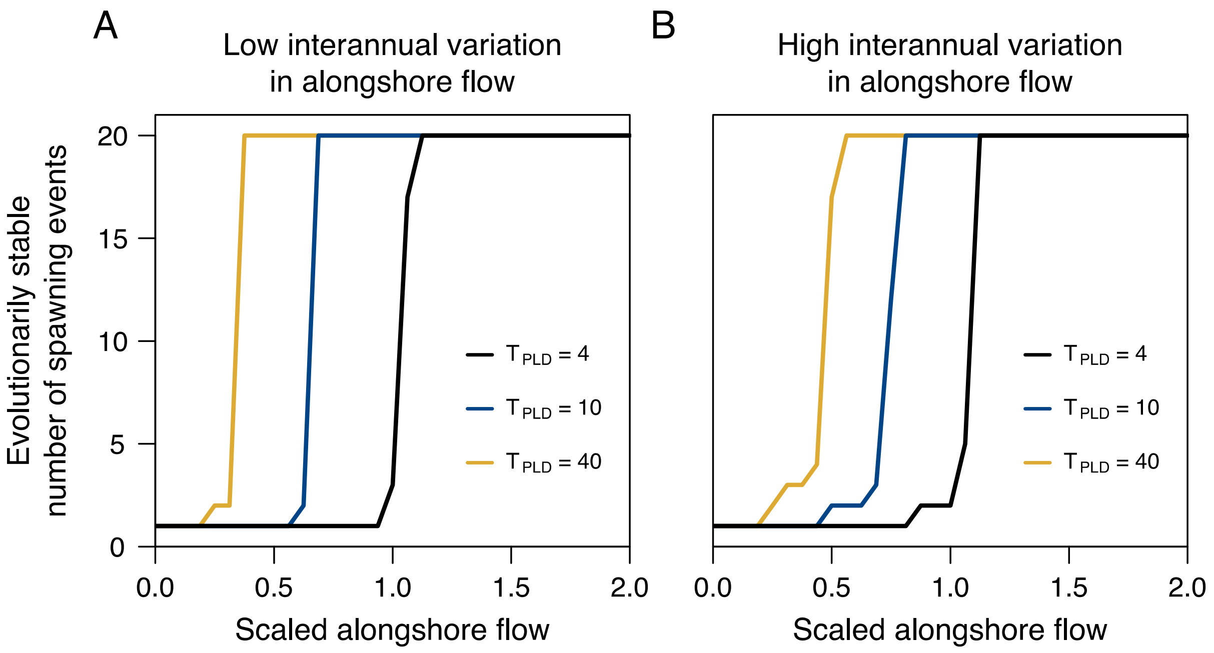


**Figure A3.** Effect of scaled alongshore flow on the evolutionarily stable $N_{spawn}$ when there is no adult mortality $A_m$=0. Panel A shows results when the standard deviation of interannual mean flow rates $\sigma_{\mathrm{IA}}$=0.012 (meters/second) and panel B shows results for when $\sigma_{\mathrm{IA}}$=0.035 (meters/second). Different colors denote different pelagic larval durations ($T_{\mathrm{PLD}}$) as indicated in the figure legend. All other parameters are the same as in figure 2.

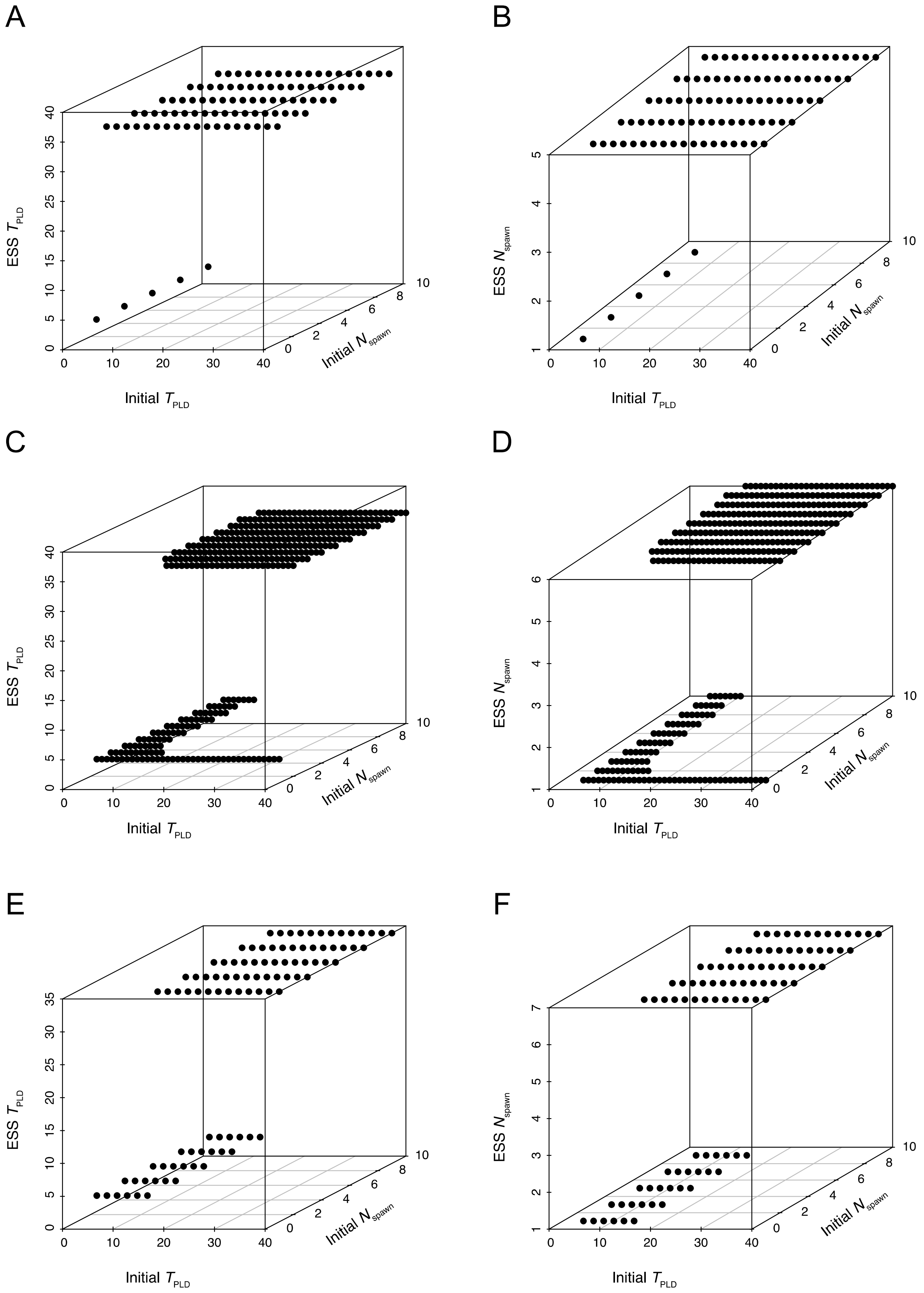
A
ESS $T_{PLD}$
Initial $T_{PLD}$
Initial $N_{spawn}$
B
ESS $N_{spawn}$
Initial $T_{PLD}$
Initial $N_{spawn}$
C
ESS $T_{PLD}$
Initial $T_{PLD}$
Initial $N_{spawn}$
D
ESS $N_{spawn}$
Initial $T_{PLD}$
Initial $N_{spawn}$
E
ESS $T_{PLD}$
Initial $T_{PLD}$
Initial $N_{spawn}$
F
ESS $N_{spawn}$
Initial $T_{PLD}$
Initial $N_{spawn}$

**Figure A4.** Evolutionarily stable values of pelagic larval durations ($T_{PLD}$) and number of spawning events ($N_{spawn}$). The first row shows results when both $T_{PLD}$ and $N_{spawn}$ evolve at the same rate. The second row shows results when $T_{PLD}$ evolves faster than $N_{spawn}$. The third row shows results when $N_{spawn}$ evolves faster than $T_{PLD}$. Parameters correspond to Figure 5B with a scaled alongshore flow rate of 1.25.

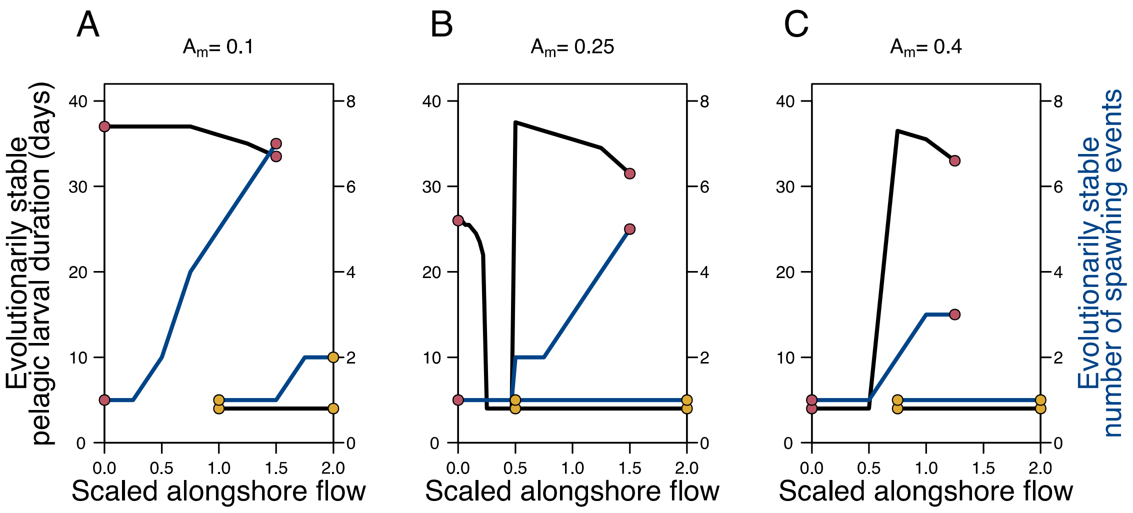


**Figure A5.** Effect of changing adult mortality rate $A_m$ on results of the model with coevolution. The evolutionarily stable values of $T_{PLD}$ (black lines) and $N_{spawn}$ (blue lines) for different degrees of scaled alongshore flow with a high standard deviation of interannual mean flow rates ($\sigma_{IA}$=0.035 meters/second). Populations always evolved to the maximum possible trait value of $N_{spawn}$, which was set to be 10. Parameters: $\sigma$=8000 (meters/day), $\tau$=4, C=0.1, $s_{crit}$=0.0103, $g$ =

0.16, $K$=200, $m$=5×10$^{-7}$, length of coastline = 100 kilometers, minimum value of $T_{PLD}$ was set to $\tau$.

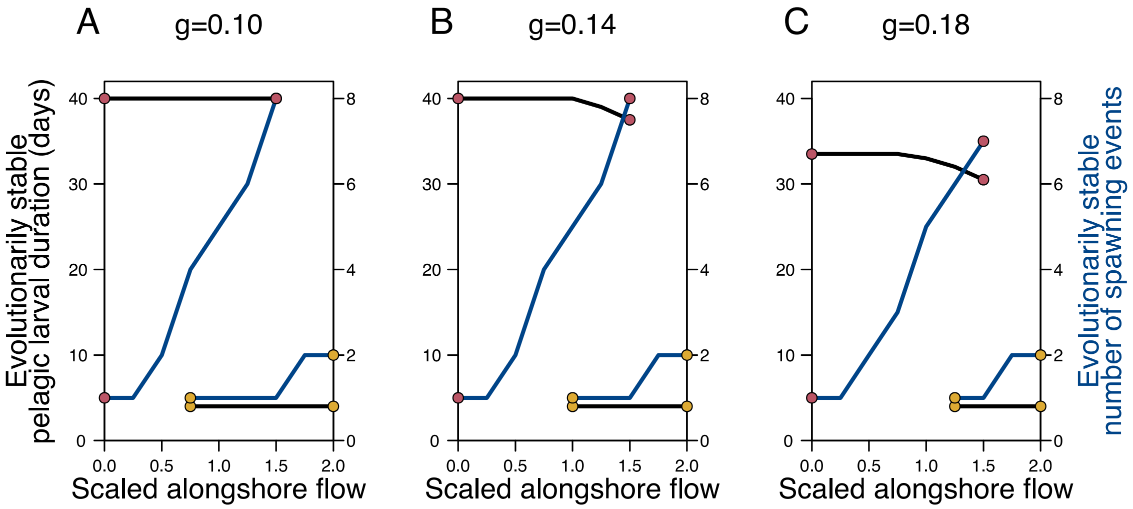


**Figure A6.** Effect of changing larval growth rate $g$ on results of the model with coevolution. The evolutionarily stable values of $T_{PLD}$ (black lines) and $N_{spawn}$ (blue lines) for different degrees of scaled alongshore flow with a high standard deviation of interannual mean flow rates ($\sigma_{IA}$=0.035 meters/second). Populations always evolved to the maximum possible trait value of $N_{spawn}$, which was set to be 10. Parameters: $\sigma$=8000 (meters/day), $\tau$=4, C=0.1, $s_{crit}$=0.0103, $K$=200, $m$=5×10$^{-7}$, $A_m$=0.1, length of coastline = 100 kilometers, minimum value of $T_{PLD}$ was set to $\tau$.

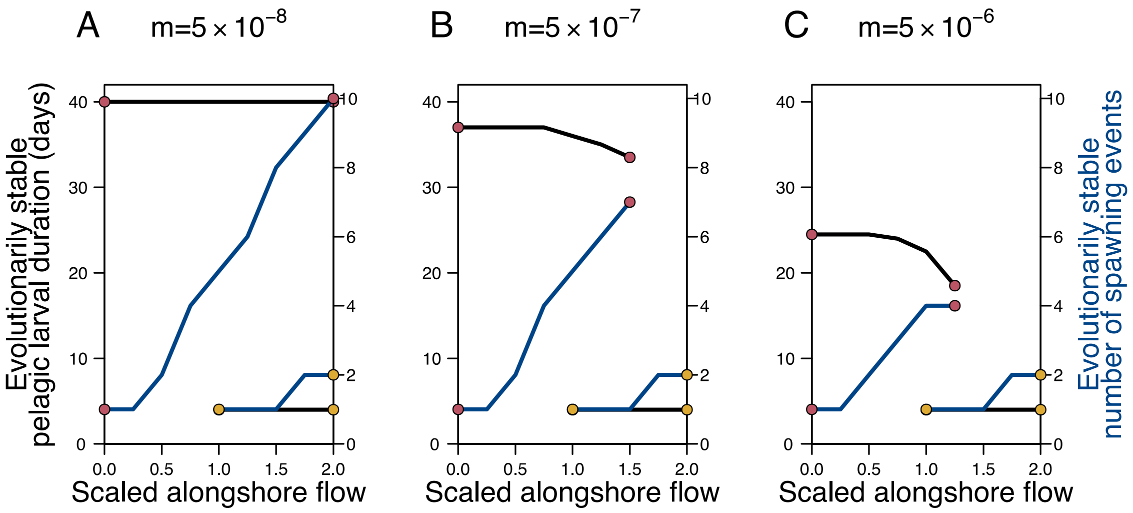


**Figure A7.** Effect of changing larval mortality rate *m* on results of the model with coevolution. The evolutionarily stable values of $T_{PLD}$ (black lines) and $N_{spawn}$ (blue lines) for different degrees of scaled alongshore flow with a high standard deviation of interannual mean flow rates ($\sigma_{IA}$=0.035 meters/second). Populations always evolved to the maximum possible trait value of $N_{spawn}$, which was set to be 10. Parameters: $\sigma$=8000 (meters/day), $\tau$=4, C=0.1, $s_{crit}$=0.0103, $K$=200, $g$=0.16, $A_m$=0.1, length of coastline = 100 kilometers, minimum value of $T_{PLD}$ was set to $\tau$.

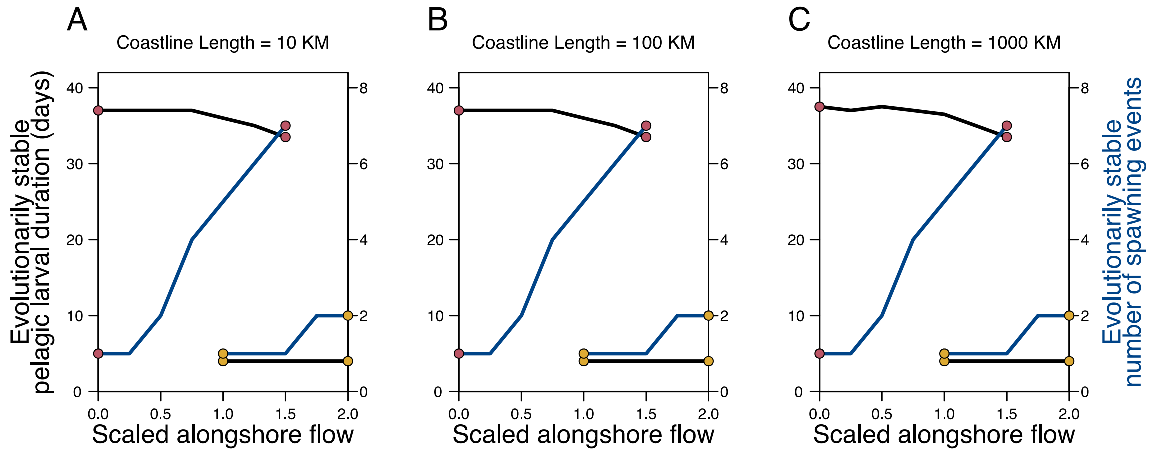


**Figure A8.** Effect of changing coastline length on results of the model with coevolution. The evolutionarily stable values of $T_{\mathrm{PLD}}$ (black lines) and $N_{\mathrm{spawn}}$ (blue lines) for different degrees of scaled alongshore flow with a high standard deviation of interannual mean flow rates ($\sigma_{\mathrm{IA}}$=0.035 meters/second). Populations always evolved to the maximum possible trait value of $N_{\mathrm{spawn}}$, which was set to be 10. Parameters: $\sigma$=8000 (meters/day), $\tau$=4, C=0.1, $s_{\mathrm{crit}}$=0.0103, $g$ = 0.16, $K$=200, $m$=5×10$^{-7}$, $A_m$=0.1, length of coastline = 100 kilometers, minimum value of $T_{\mathrm{PLD}}$ was set to $\tau$.

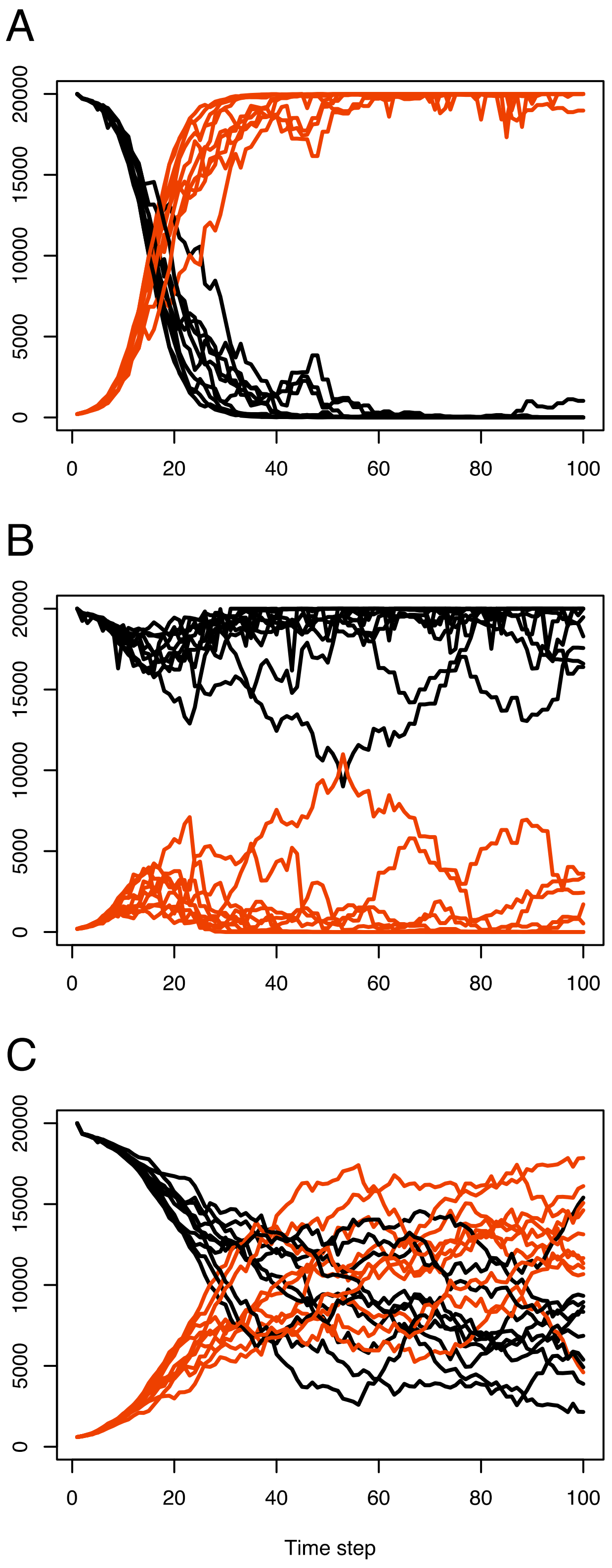
A
B
C
0
5000
10000
15000
20000
0
20
40
60
80
100
Time step

**Figure A9.** Evolution of pelagic larval duration ($T_{PLD}$) in the stochastic simulations. Each panel shows the results of 10 runs of the simulation. Black lines denote the population size of the resident ($T_{PLD}$ = 30) and red lines denote the population size of the mutant ($T_{PLD}$ = 25). In Panel A, scaled advection = 0.25 and the number of spawning events ($N_{spawn}$) is 1. Note that in this case the mutant typically successfully invades. In Panel B, scaled advection = 1.0 and $N_{spawn}$ = 1. Note that in this case the mutant typically does not successfully invades. In Panel C, scaled advection = 1.0 and $N_{spawn}$ = 5. Note that in this case the mutant typically successfully invades. Parameters: $\sigma$=4000 (meters/day), $\tau$=4, C=0.1, $s_{crit}$=0.0103, $g$ = 0.16, $m$ =$5\times10^{-7}$, $A_m$=0.0, $K$=200, length of coastline = 100 kilometers.

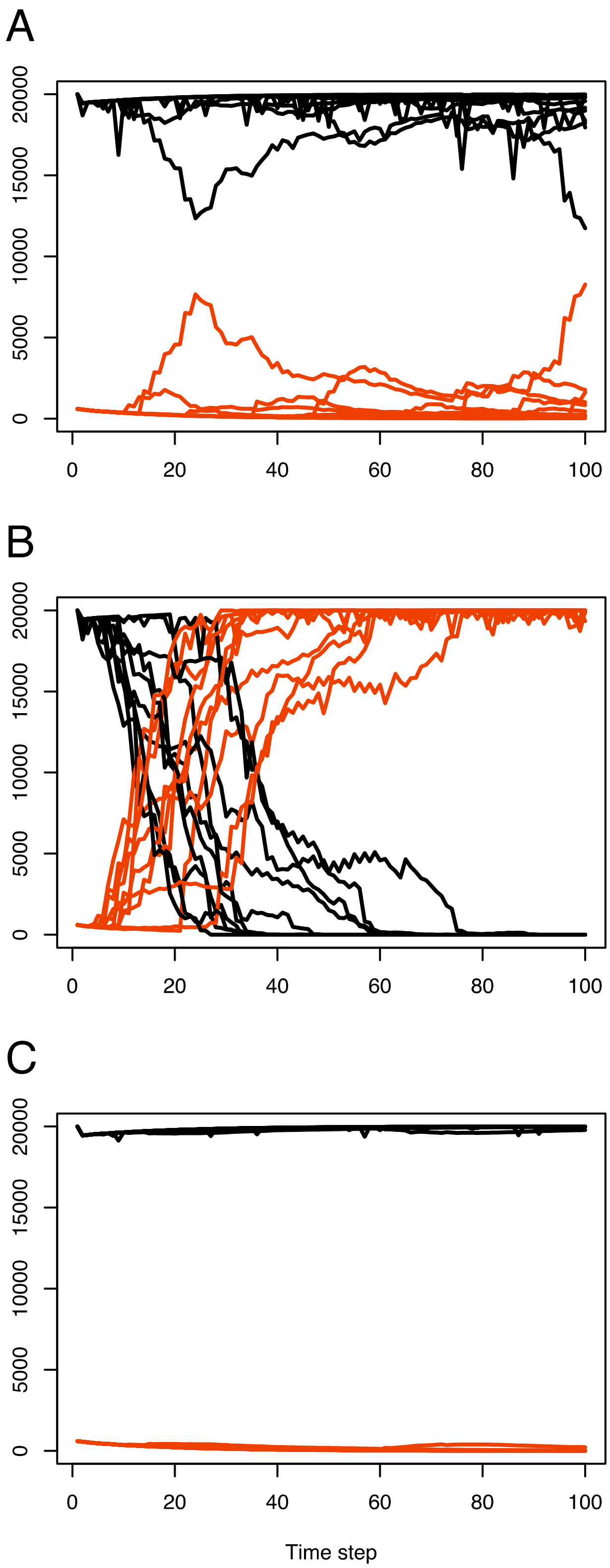
A
B
C
20000
15000
10000
5000
0
0
20
40
60
80
100
Time step

**Figure A10.** Evolution of the number of spawning events ($N_{\mathrm{spawn}}$) in the stochastic simulations. Each panel shows the results of 10 runs of the simulation. Black lines denote the population size of the resident ($N_{\mathrm{spawn}}$ = 1) and red lines denote the population size of the mutant ($N_{\mathrm{spawn}}$ = 5). In Panel A, scaled advection = 0.25 and the pelagic larval duration ($T_{\mathrm{PLD}}$) is 40. Note that in this case the mutant typically does not successfully invades. In Panel B, scaled advection = 1.0 and $T_{\mathrm{PLD}}$ = 10. Note that in this case the mutant typically successfully invades. In Panel C, scaled advection = 1.0 and $T_{\mathrm{PLD}}$ = 40. Note that in this case the mutant typically does not successfully invades. Parameters: $\sigma$=4000 (meters/day), $\tau$=4, C=0.1, $s_{\mathrm{crit}}$=0.0103, $g$ = 0.16, $m$ =$5\times10^{-7}$, $A_m$=0.0, $K$=200, length of coastline = 100 kilometers.
